# Intragenomic conflict underlies extreme phenotypic plasticity in queen-worker caste determination in honey bees (*Apis mellifera*)

**DOI:** 10.1101/2024.06.09.598129

**Authors:** Sean T. Bresnahan, Shaun Mahony, Kate Anton, Brock Harpur, Christina M. Grozinger

## Abstract

Caste determination of honey bees (*Apis mellifera*) is a prime example of developmental plasticity, where differences in larval diet will result in identical genotypes yielding either long-lived, reproductive queens or short-lived, facultatively sterile workers. Beyond environmental factors, intragenomic conflict between genes inherited from the mother (matrigenes) versus the father (patrigenes) is also hypothesized to generate this plasticity. In honey bees, the Kinship Theory of Intragenomic Conflict predicts selection on patrigenes to enhance traits that result in fitness gained through reproduction, and thus patrigenes should favor the queen caste fate. Here, we conducted allele-specific transcriptome analyses on queen-destined larvae (QL) and worker-destined larvae (WL) at 192 hours post-fertilization (hpf), a critical stage for caste determination. Our findings reveal hundreds of genes with parent-of-origin effects (POEs), with significant patrigene-biased transcription in QL. Genes with POEs in honey bees resemble imprinted genes in other taxa in terms of genomic clustering, recombination rate, intron length and CpG density, and a subset are maintained from 24hpf eggs. Previous studies demonstrated that DNA methylation, the canonical regulatory mechanism underlying transcriptional POEs in placental mammals, angiosperms, and some insects, is not operating in honey bees or other social insects. We use allele-specific ChIP-seq analyses to demonstrate that POEs on caste-specific histone post-translational modification (HPTM) profiles of H3K27me3, H3K4me3 and H3K27ac are associated with POEs on transcription. Together, these findings suggest that parent-of-origin intragenomic conflicts may contribute broadly to phenotypic plasticity and may be associated with HPTMs, suggesting a “non-canonical” genomic imprinting-like system in social insects.

## Background

Eusocial insects (which including certain species of bees, wasps, ants, and termites) present exceptional models for studies of developmental plasticity (Yoon *et al*., 2023), where a single genotype yields a spectrum of phenotypes in response to environmental variation (Pigliucci, 2001). In these species, large self-organizing colonies are formed of individuals partitioned into reproductive and typically nonreproductive phenotypes known as castes: two developmental fates of totipotent larvae that are canalized by environmental differences–typically in nutrition–experienced during a critical early growth window (Thompson & Chernyshova, 2011). The two castes represent organisms with distinct repertoires of morphological, physiological, and behavioral characteristics, which cooperate to maximize colony fitness. Beyond environmental factors, intragenomic conflict between genes inherited from the mother (matrigenes) and father (patrigenes) is also hypothesized to generate this plasticity (Queller, 2003). In this study, we examined whether eusocial insect caste determination may be influenced by parent-of-origin intragenomic conflict.

Gene regulatory mechanisms play a central role in developmental plasticity, enabling context-specific control over gene expression. Genomic imprinting – a regulatory process that results in parent-of-origin gene expression – is an adaptive model of developmental plasticity, providing a mechanism for balancing competing interests of matrigenes and patrigenes within organisms, particularly over reproduction (Radford et al., 2011). In Eutherian mammals, genomic imprinting in the placenta plays a crucial role in balancing competing interests between the mother and father by influencing maternal resource allocation, and by regulating placental growth and maternal immunological responses to invasion of the placental trophoblast (Hanna, 2020). In angiosperms, imprinting in the endosperm mediates similar conflicts over nutrient transfer and tissue proliferation (Gehring & Satyaki, 2017).

Several theories try to explain the evolution of genomic imprinting (reviewed in Patten *et al*., 2014). Here, we examine the Kinship Theory of Intragenomic Conflict (KTIC), which proposes that maternal and paternal interests are distinct in organisms where the female can bear offspring derived from multiple males (Haig, 2000). In such cases, the maternal interest is to equally allocate resources among all offspring, whereas the paternal interest is to maximize only the survival of offspring carrying the same alleles. Thus, matrigenes and patrigenes can have different strategies for maximizing their reproductive fitness, with conflicting developmental effects, reflecting the distinct interests of each parent (Gardner & Úbeda, 2017). These different selective pressures ultimately lead to parent-of-origin effects (POEs) on gene expression, with opposite transcriptional patterns for matrigenes and patrigenes.

The western honey bee *Apis mellifera* is an excellent model for studies of developmental plasticity, including parent-of-origin intragenomic conflicts and their regulatory basis (Pegoraro *et al*., 2017). Honey bee colonies have a single reproductive female queen and tens of thousands of her daughters (workers) which typically remain sterile and forego their own reproduction to rear the queen’s offspring (Page, 2013). Since honey bees are haplodiploid (females are diploid and males are haploid) and queens are polyandrous, her daughters on average share more of their matrigenes than patrigenes. Thus, matrigenes and patrigenes are unevenly distributed among nestmates and should experience different selective pressures leading to POEs on transcription (Queller, 2003). Such parent-of-origin intragenomic conflicts have been demonstrated in association with behavioral variation in adult worker honey bees (Bresnahan *et al*., 2023a, 2023b; Galbraith *et al*., 2016a, 2021; Gibson *et al*., 2015; Kocher *et al*., 2015), and in the transcriptomes of diploid eggs laid by queens (Smith *et al*., 2020).

Female queens and workers are produced from diploid eggs, but queen-destined larvae are fed a different diet (royal jelly) by workers throughout their larval development, while worker-destined larvae are fed royal jelly for the first three days after hatching and then switched to a diet of worker jelly (Alhosin, 2023). Workers will rear queens under certain circumstances, such as when the old queen is lost or failing (reviewed in Sagili *et al*., 2018). While there are clearly fitness advantages to a larva being reared as a queen, the colony must regulate the production of new queens to reduce conflicts and maximize group fitness (Ferreira *et al*., 2024); thus, intragenomic conflict may partially mediate this inter-colony conflict, with patrigenes and matrigenes promoting different developmental outcomes. According to the KTIC, patrigenes should favor the development of the reproductive queen caste over the nonreproductive worker caste (Queller, 2003). In this study, we used a reciprocal cross design (see Methods: Figure 10) to assess POEs on transcriptional profiles of honey bee worker and queen-destined larvae (WL and QL, respectively) at 192 hours post-fertilization (hpf), a point at which the caste fate is canalized and irreversible (Wojciechowski *et al*., 2018). We conducted an allele-specific transcriptome analysis to test the hypothesis that the reproductive caste fate (QL) is associated with enrichment for patrigene-biased transcription compared to the nonreproductive caste fate (WL).

Since these selective advantages and expression differences arise from the origin of the gene (maternal-or paternal-origin) and not the gene sequence, POEs provide a unique opportunity to study sequence-independent mechanisms of phenotypic variation. Consequently, parent-of-origin gene regulation is one of the most longstanding paradigms for investigating the phenotypic outcomes of interactions between genotype, phenotype, and evolution (Ferguson-Smith, 2011). In studies in vertebrates, plants, and some insects, POEs on transcription are generated by parental allele-specific DNA methyl-cytosine (DNAme) of introns, promoters, or other *cis*-elements (MacDonald, 2012; Batista & Köhler, 2020; Hanna & Kelsey, 2021). However, though POEs on transcription have been found in three social insect species, honey bees (Kocher *et al*., 2015; Smith *et al*., 2020; Galbraith *et al*., 2016, 2021), bumble bees (Marshall *et al*., 2020), and *Nasonia* jewel wasps (Olney *et al*., 2021), these are not associated with DNAme in these species (Wu *et al*., 2020; Marshall *et al*., 2020; Olney *et al*., 2021). Other studies have demonstrated that DNAme is not associated with downregulating or silencing gene expression in social insects (Maleszka & Kucharski, 2022), but rather may contribute to mediating transcriptional noise (Wu *et al*., 2020; Bogan & Yi, 2024). Thus, how parent-of-origin transcription is regulated in social insects remains an open area of investigation (Pegoraro *et al*., 2017; Oldroyd & Yagound, 2021), and some understanding of these mechanisms is necessary to direct investigations into the nature of their establishment and inheritance.

Across taxa, various histone 3 (H3) post-translational modifications (HPTMs) which promote or restrict transcription have recently been shown to be essential for POEs on certain genes in the absence of DNAme (Soliman & Coughlan, 2024). In placental mammals and angiosperms, DNA methylation asymmetries between female and male genomes are caused by extensive demethylation in female, but not male, germ cells. This DNA demethylation in female germ cells enables recruitment of DNA-binding and Polycomb group (PcG) proteins that catalyze trimethylation of lysine 27 of H3 (H3K27me3). In these species, maternally-inherited H3K27me3 at some loci promotes maternal allele-specific chromatin condensation, silencing the maternal allele and ensuring only the paternal allele is accessible for transcriptional activity (Batista & Köhler, 2020; Hanna & Kelsey, 2021; Raas *et al*., 2021). In *Drosophila* – which do not possess the canonical DNA methyltransferases (Krauss & Reuter, 2011) – parental allele-specific transcriptional variation has been associated with parental allele-specific variation in promoter activity, indicated by trimethylation of lysine 4 of H3 (H3K4me3), and enhancer activity, indicated by acetylation of lysine 27 of H3 (H3K27ac) (Floc’hlay *et al*., 2021). Thus, POEs on chromatin regulation may function as an alternative gene regulatory mechanism of parent-of-origin intragenomic conflicts in species which do not use DNAme for transcriptional silencing (Inoue, 2023).

Although previous research demonstrates that honey bee WL and QL show caste-specific differences in genome-wide enrichment of H3K27ac and H3K4me3 (Wojciechowski et al., 2018), and adult worker bees show variation in genome-wide enrichment of H3K27me3 during ovarian development (Duncan et al., 2020), whether these HPTMs contribute to POEs on transcription in this species is unknown. In this study, we assessed variation in genome-wide enrichment profiles of, and POEs on, H3K27me3, H3K27ac and H3K4me3 in QL and WL at 192hpf, to test the hypothesis that POEs on transcription are associated with POEs on HPTM enrichment. We conducted an allele-specific ChIP-seq analysis to test the hypothesis that the reproductive caste fate (QL) is associated with enrichment for patrigene-biased HPTMs indicative of active transcription (H3K4me3 and H3K27ac) compared to the nonreproductive caste fate (WL). Additionally, we tested the hypothesis that genes showing POEs on transcription are associated with genomic regions showing POEs on HPTM enrichment, with H3K4me3 and H3K27ac associated with more expressed alleles, and H3K27me3 associated with silenced alleles.

These studies also provided an opportunity to examine additional questions of how genes showing POEs on transcription might contribute to phenotypic variation, whether they are maintained across developmental stages, and whether they exhibit traits like those of imprinted genes in other taxa. Previous studies in honey bees found that genes showing POEs on transcription do not typically show differences in gene dosage between phenotypes (Bresnahan *et al*., 2023a, 2023b; Galbraith *et al*., 2016a, 2021), presumably because of selection for a balance on the individual effects of matrigenes and patrigenes engaged in conflict (Haig, 2008; Patten *et al*., 2016). Instead, like imprinted genes (Al Adhami *et al*., 2015; Macias-Valesco *et al*., 2022), genes with POEs in honey bees are frequently enriched in co-expressed gene networks (Bresnahan *et al*., 2023b; Galbraith *et al*., 2021), suggesting they may contribute to phenotypic variation through cascading effects on e.g., cell signaling pathways. In this study we examined relationships between POEs on transcription, differential gene expression and gene networks, to further explore how genes engaged in parent-of-origin intragenomic conflicts may contribute to caste determination.

It is unclear whether genes with POEs in social insects are truly “imprinted”, or whether parent-of-origin intragenomic conflict represents a distinct gene regulatory mechanism of phenotypic plasticity (Oldroyd & Yagound, 2021; Pegoraro *et al*., 2017). Hallmarks of imprinted genes include maintenance of POEs on transcription across cell divisions (Sanli & Feil, 2015), clustering throughout the genome (as imprinted genes are frequently regulated by shared enhancers or imprinting control domains) (Wolf, 2013), relatively higher recombination rates (Sandovici *et al*., 2006), and longer, GC- and CpG-rich introns compared to unbiased genes (Hutter *et al*., 2010). In this study, we explored whether genes with POEs in 24hpf eggs maintain their parental allele-specific expression status in larvae. Additionally, we examined whether genes with POEs are clustered throughout the honey bee genome, have longer, GC- and CpG-rich introns, and exhibit higher rates of recombination within *Apis* compared to unbiased genes in these tissues.

Here, we investigate the extreme phenotypic plasticity of queen-worker caste determination in honey bees to test predictions of the KTIC, evaluate whether a non-canonical genomic imprinting-like mechanism is associated with POEs on gene expression, and determine if genes showing POEs exhibit genomic and sequence features found in imprinted genes characterized in other taxa, which would suggest these genes experience similar selective processes over evolutionary time. To conduct these studies, we used instrumental insemination to create reciprocal crosses between honey bees derived from different genetic stocks in two distinct genetic Blocks and manipulated the larvae resulting from these crosses to induce development of either the worker or queen caste fate.

## Results

### Transcriptomic differences underpinning reproductive caste determination

We evaluated transcriptomes of *n*=10 queen-destined larvae (QL) and *n*=10 worker-destined larvae (WL) for each of two reciprocal crosses within two distinct genetic Blocks (*n*=40 larvae in total; Blocks 1 and 2, see Methods). Like previous studies (Barchuck *et al*., 2007; Ashby *et al*., 2016; Wojciechowki *et al*., 2018; Wang *et al*., 2021; He *et al*., 2022; Zhang *et al*., 2023), we identified hundreds of differentially expressed genes (DEGs) between QL and WL at 192 hours post-fertilization (hpf) (Figure 1a, Supplementary Materials: S2.1). *K*-means clustering of the transcriptomes with *k*=2 using either all *n*=11,705 expressed genes (Figure 1b) or the *n*=511 DEGs (Figure 1c) shows that the two castes are transcriptionally distinct phenotypes at 192hpf. DEGs were overrepresented among KEGG pathways for hormone biosynthesis, tyrosine metabolism, and metabolic pathways, genes encoding signaling peptides and proteins involved in olfaction and sensory transduction (Supplementary Materials: S8.1.

**Figure 1.**
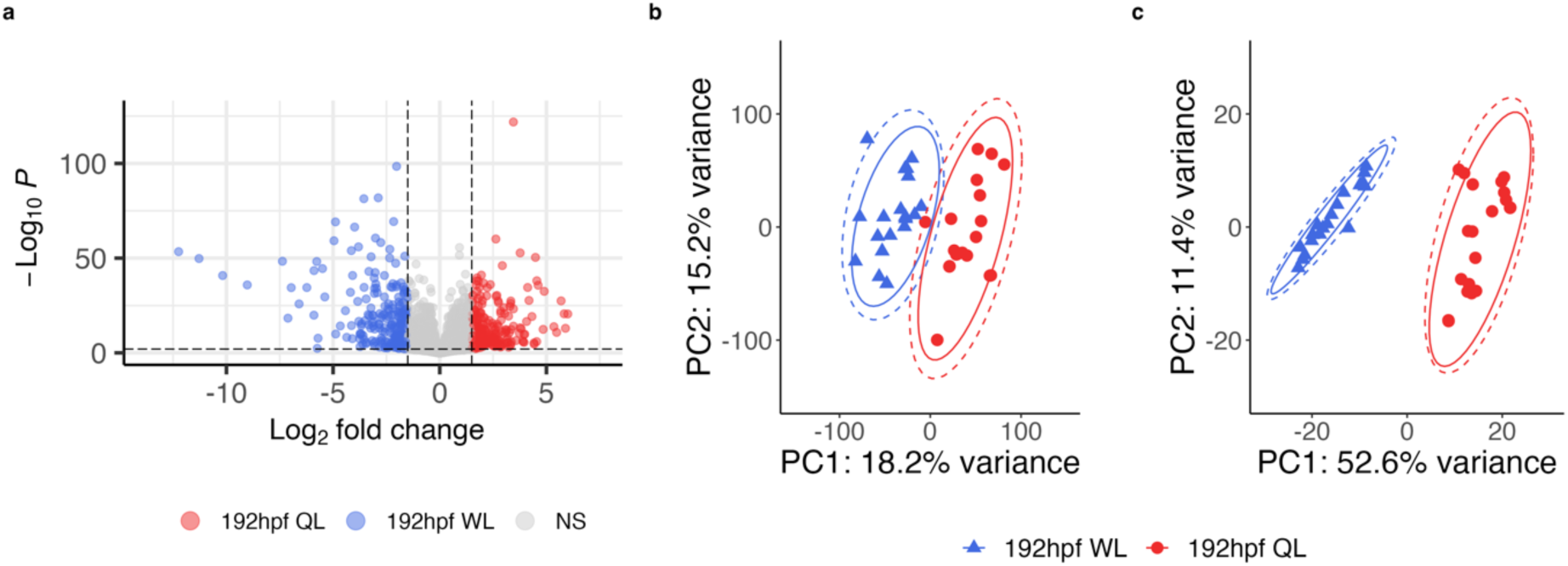
Transcriptomic differences between worker-destined larvae (WL) and queen-destined larvae (QL) at 192hpf. **(a)** Volcano plot of the *n*=11,705 genes expressed in QL and WL. Dashed lines indicate thresholds of |log_2_ fold change (FC)| >1.5 and FDR <0.01. Colors indicate whether a gene was significantly upregulated in QL (red), upregulated in WL (blue), or showed no significant difference in expression between castes (NS, grey). **(b-c)** *K*-means clustering on *k*=2 of **(b)** all *n*=11,705 expressed genes and **(c)** the *n*=511 differentially expressed gene.

### Parent-of-origin effects (POEs) on transcription patterns underlying reproductive caste determination support predictions of the Kinship Theory of Intragenomic Conflict

The genomes of the mothers (queens) and fathers (drones) of the reciprocal crosses in each Block were sequenced (*n*=4 queens and *n*=4 drones in total). At approximately 87× genome coverage, we detected 407,300 (±28,335) high-confidence homozygous SNPs per F1 diploid queen and 887,609 (±162,105) per F1 haploid drone. Of these SNPs, 252,179 (±46,961) were within transcripts and varied in their sequence identity between the parents of each cross, allowing for the identification of parent-of-origin reads in the F2 larvae. The number of SNPs and transcripts examined in this study after filtering SNP positions with low mRNA-seq read coverage in the F2 larvae, and number of transcripts showing parent-of-origin effects (POEs) on transcription in each Block are listed in Supplementary Materials: S3.1-3.2. In support of our hypothesis, we found that patrigene-biased transcripts were enriched in QL relative to WL in both Blocks (see Figure 2 for results from Block 2 and Supplementary Materials: S3.3 for results from Block 1). Overall, transcripts showing POEs in larvae were enriched for genes encoding developmental proteins, and genes related to axon guidance and synaptic target recognition (Supplementary Materials: S8.1).

**Figure 2.**
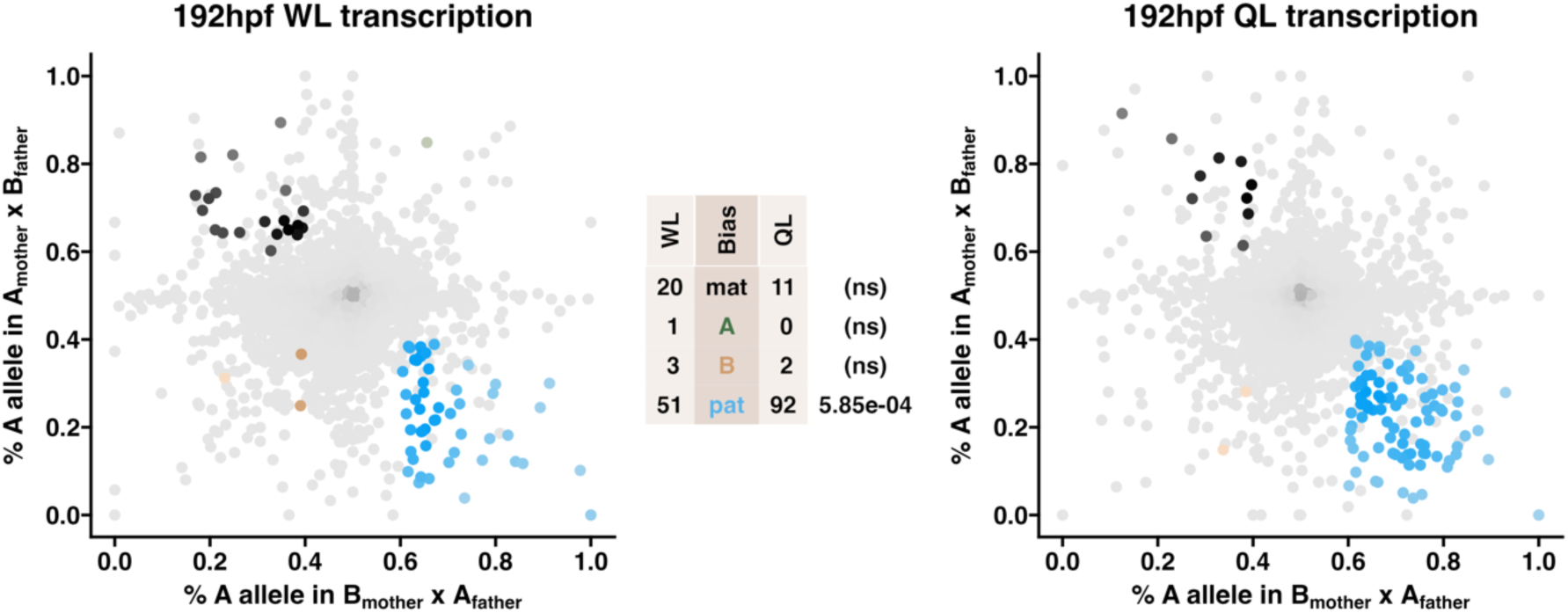
Queen-destined larvae show enriched patrigene-biased transcription relative to worker-destined larvae. Allele-specific transcriptomes were assessed in F2 worker-destined larvae (WL) and queen-destined larvae (QL) at 192hpf collected from a reciprocal cross between different stocks of European honey bees (See Methods: Figure 10, Block 2). The x-axis represents, for each transcript, the proportion of lineage A reads in bees with a lineage B mother and lineage A father (*p1*). The y-axis represents, for each transcript, the proportion of lineage A reads in bees with a lineage A mother and lineage B father (*p2*). Each color represents a transcript which is significantly biased at all tested SNP positions: black is maternal (mat), green is lineage A, gold is lineage B, blue is paternal (pat), and grey is not significant. Center table: the number of transcripts showing each category of allelic bias and *p*-values for Chi-squared tests of independence for comparisons between the castes are indicated (NS = not significant). Significance of allele-biased transcription was determined using the overlap between two statistical tests: a general linear mixed model (GLIMMIX), and a Storer-Kim binomial exact test along with previously established thresholds of *p1*<0.4 and *p2*>0.6 for maternal bias, *p1*>0.6 and *p2*<0.4 for paternal bias, *p1*<0.4 and *p2*<0.4 for lineage B bias and *p1*>0.6 and *p2*>0.6 for lineage A bias (Wang & Clark, 2014). Similar results were obtained in Block 1 (Supplementary Materials: S3.3).

### Parent-of-origin intragenomic conflicts may contribute to caste determination through differential gene expression and gene networks

We compared the set of differentially expressed genes (*n*=511) and genes with POEs (*n*=508) in both Blocks and found that these two sets overlap (*n*=55) more than expected by chance (*X^2^*-test, *p*=2.2×10^-16^). Genes that were both differentially expressed and showed POEs were overrepresented for metabolic pathways and signaling peptides, including *vitellogenin* and *major royal jelly protein 1* (Supplementary Materials: S8.1). Thus, parent-of-origin intragenomic conflict may contribute to caste determination by altering gene dosage of core genes involved in caste determination.

As in previous studies (Galbraith *et al*., 2020; Bresnahan *et al*., 2023b), we also tested the hypothesis that genes engaged in parent-of-origin intragenomic conflicts should be enriched in gene networks (Patten *et al*., 2016). Using a Weighted Gene Co-Expression Network Analysis, we identified seventeen co-expressed gene modules consisting of *n*=524 (±463) genes per module. Ten modules were significantly correlated with caste, two of which (Modules 2 and 4) were significantly enriched with genes showing POEs on transcription (Supplementary Materials: S2.2). Module 2 consisted of *n*=1,717 genes, *n*=127 of which showed POEs on transcription (Fisher’s exact test, *p*=0.02), had the highest positive correlation with caste (*r*=0.85, *p*=1×10^-11^), and was enriched for genes in KEGG pathways for carbon metabolism, glycolysis/gluconeogenesis, and genes related to transmembrane transport (Supplementary Materials: S8.2). Module 4 consisted of *n*=846 genes, *n*=70 of which showed POEs (Fisher’s exact test, *p*=0.007), was positively correlated with caste (*r*=0.54, *p*=1×10^-4^), and was enriched for genes in KEGG pathways for nucleocytoplasmic transport, spliceosome, RNA polymerase, and ribosome biogenesis, in addition to several GO terms for chromatin remodeling, mRNA splicing via spliceosome, and transcription by RNA polymerase I (Supplementary Materials: S8.2). Thus, parent-of-origin intragenomic conflicts may also contribute to caste determination through cascading effects on developmental signaling pathways and on gene networks related to transcriptional regulation.

### Transcription dynamics associated with histone post-translational modifications (HPTMs)

We determined and compared between castes the genome-wide distribution of three HPTMs with functions in the regulation of transcription and *cis*-elements: H3K27me3, H3K27ac, and H3K4me3 (Bannister & Kouzarides, 2011). For each HPTM, we identified thousands of peaks of enrichment genome-wide in both castes (Table 1, Supplementary Materials: S5.1). Like in other taxa, we find that all three HPTMs are enriched at transcription start sites (TSS) of transcribed genes (Supplementary Materials: S4.1).

**Table 1.** Frequency table of consensus peaks (identified in both crosses within a Block).

| HPTM | Caste | Peaks |
| --- | --- | --- |
| H3K27me3 | WL | 41,742 ($\pm 1,120$) |
| | QL | 45,954 ($\pm 14,465$) |
| H3K27ac | WL | 36,523 ($\pm 12,848$) |
| | QL | 45,960 ($\pm 12,572$) |
| H3K4me3 | WL | 44,434 ($\pm 8,496$) |
| | QL | 50,718 ($\pm 12,544$) |

Mean count of peaks in Blocks 1 and 2 and standard deviations between Blocks are indicated. Peaks of enrichment were identified using MACS2 and relative to input samples. See Supplementary Materials: S5.1 for full details.

As the relationship between each HPTM and transcription has been shown to vary between gene bodies and promoters (Bowman & Poirier, 2015), we fit logistic regression models to examine relationships between transcript abundance and peak presence/absence across these features for each caste and Block, separately (Figure 3; Supplementary Materials: S6.1-6.3). H3K27me3 is a dynamic mark, typically indicative of transcriptional silencing, but also active transcription in certain contexts (McEwen & Ferguson-Smith, 2010; Young *et al*., 2011). We found a significant negative association between transcript abundance and H3K27me3 enrichment in promoters (Figure 3a), but no association in gene bodies (Figure 3d). In contrast, H3K27ac and H3K4me3 are typically indicative of active transcription (Beacon *et al*., 2021). We found significant positive associations between transcript abundance and both H3K27ac and H3K4me3 enrichment in gene bodies (Figure 3e-f), but no association between transcript abundance and either mark in promoters (Figure 3b-c). Randomly generated sets of genomic intervals of the same quantity and average length of the peaks in each sample group did not show the same associations as the true peak sets (Supplementary Materials: S6.4).

**Figure 3.**
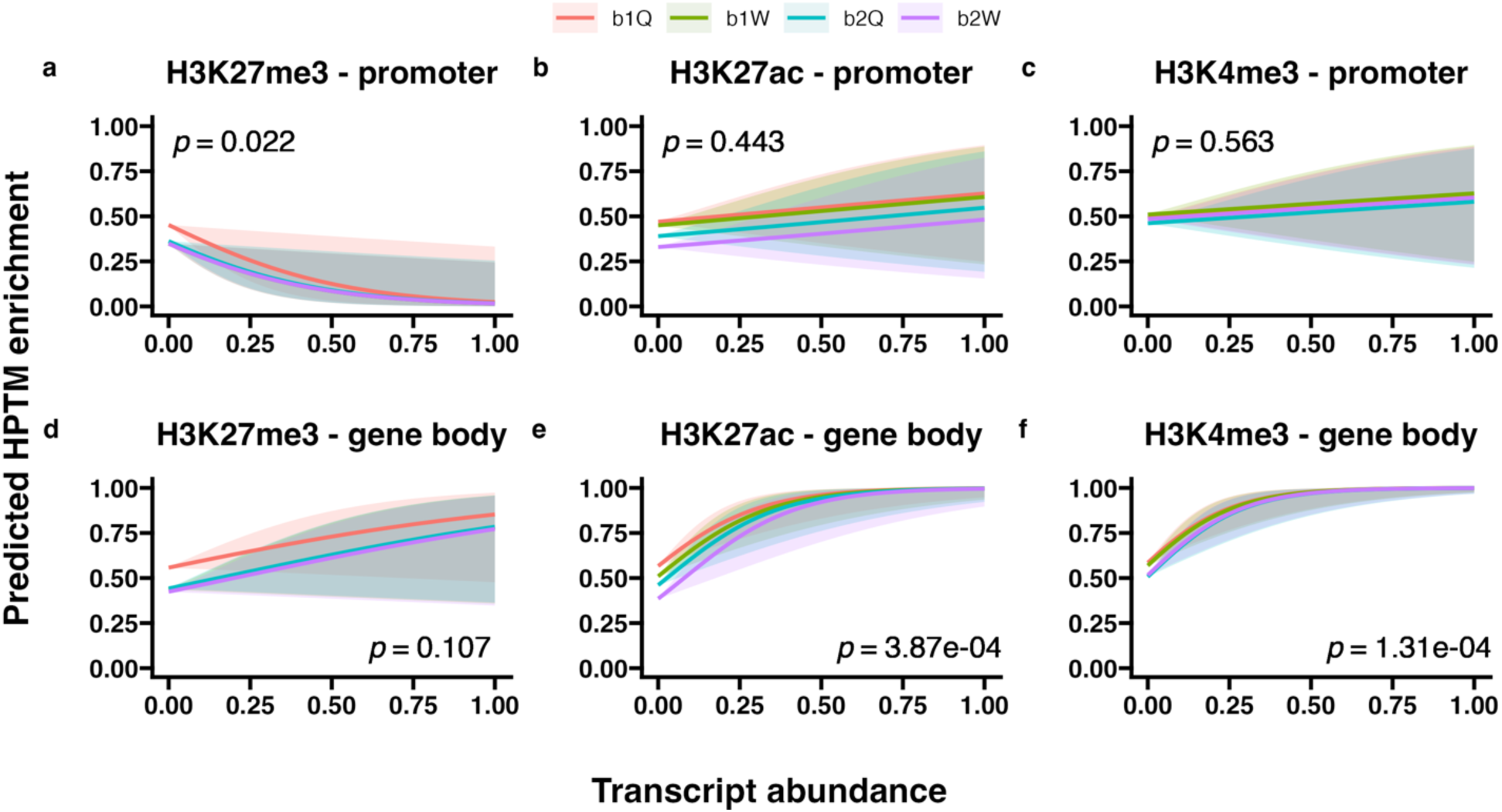
Predicted enrichment of each HPTM across **(a-c)** promoters and **(d-f)** gene bodies given transcript abundance in each caste and Block. 95% confidence intervals are indicated. Transcript abundance estimates were pooled across samples for each Block (b1 & b2) and caste (Q & W), separately, normalized using the variance stabilizing transformation, and standardized to a range of 0<*x*<1. For the response variable, genes were scored as “1” if a peak overlapped the gene or “0” if no peak overlapped the gene. See Supplementary Materials: S6 for associated odds ratios tests and comparisons to randomly generated peak sets.

### HPTM enrichment differences underpin reproductive caste determination

We found that the castes vary in distribution of H3K27me3, with 187/30,778 (∼0.6%) regions showing differential enrichment (Figure 4a, Supplementary Materials: S4.2). Like Wojciechowski *et al*. (2018), we found that 192hpf WL and QL vary in distribution of H3K27ac, with 36/31,993 (∼0.11%) regions being differentially enriched (Figure 4b), and of H3K4me3, with 748/32,518 (∼2.3%) differentially enriched regions (Figure 4c). Genes within differentially enriched H3K27me3 regions were overrepresented for transducer and receptor proteins. Within differentially enriched H3K4me3 regions, genes were overrepresented for proteins related to transcription regulation and mRNA surveillance. No GO terms or KEGG pathways were overrepresented among genes within differentially enriched H3K27ac regions (Supplementary Materials: S8.3).

**Figure 4.**
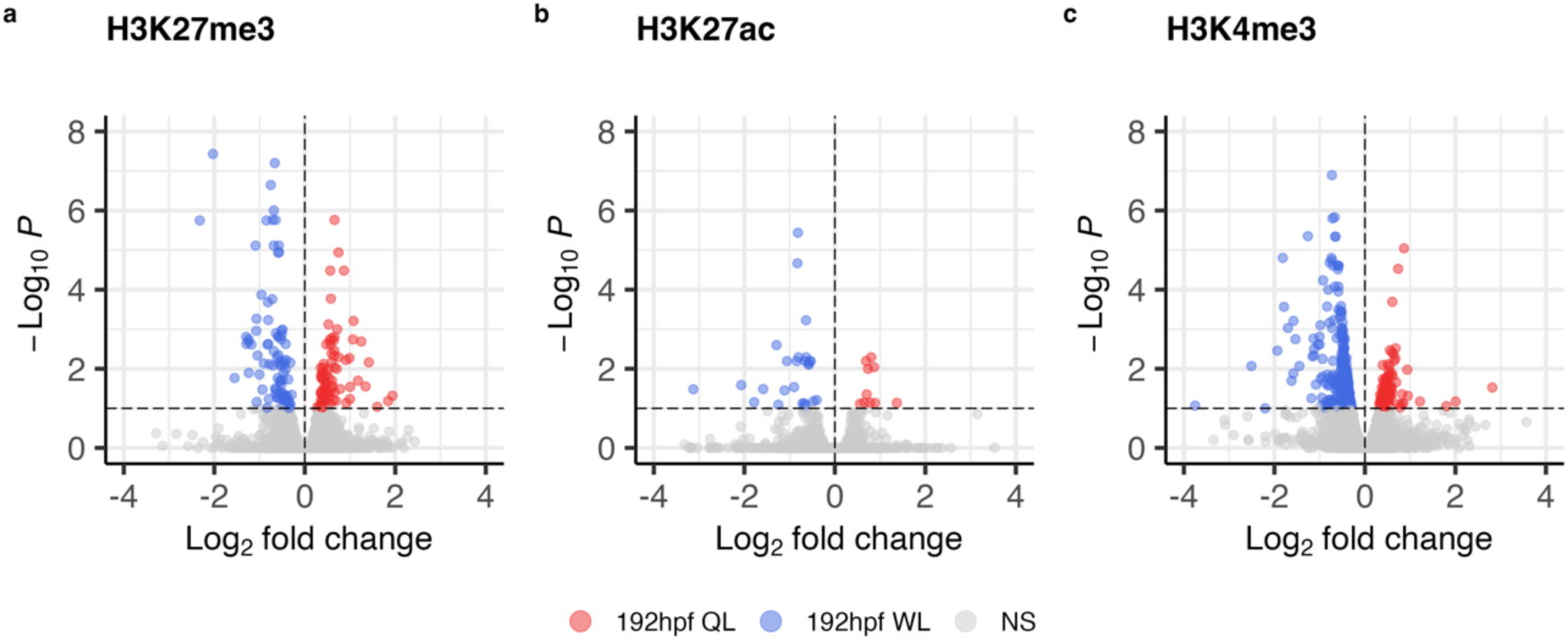
Differences in enrichment of HPTMs between queen-destined larvae (QL) and worker-destined larvae (WL) at 192hpf, for **(a)** *n*=30,788 regions for H3K27me3 **(b)** *n*=31,993 regions for H3K27ac, and **(c)** *n*=32,518 regions for H3K4me3. Dashed lines indicate thresholds of |log_2_(FC)| >0 and FDR <0.1. Colors indicate whether a window showed higher enrichment in QL (red), WL (blue), or showed no significant difference in enrichment between the castes (NS, grey).

### POEs on HPTM enrichment in caste determination

For each HPTM, we identified hundreds of peaks showing POEs on enrichment in both castes (Figure 5, Supplementary Materials: S5). We found that QL showed more patrigene-biased enrichment of all three HPTMs compared to WL across both promoters and gene bodies in both Blocks, and more matrigene-biased enrichment of H3K27me3 in gene bodies in Block 2. In contrast, WL showed more matrigene-biased H3K4me3 enrichment compared to QL across gene bodies in both Blocks and across promoters in Block 2, and more matrigene-biased H3K27ac enrichment across gene bodies and promoters in Block 1. Thus, paternal-origin chromatin in QL shows more transcriptional and regulatory activity compared to WL. Moreover, maternal-origin chromatin in QL is marked for more regulatory activity compared to WL, but this result is influenced by colony genotype. In contrast, maternal-origin chromatin in WL is marked for more transcriptional activity compared to QL, but this result is also influenced by colony genotype. See Supplementary Materials: S5 for full details of POEs on enrichment of each HPTM in each Block separated by gene region.

**Figure 5.**
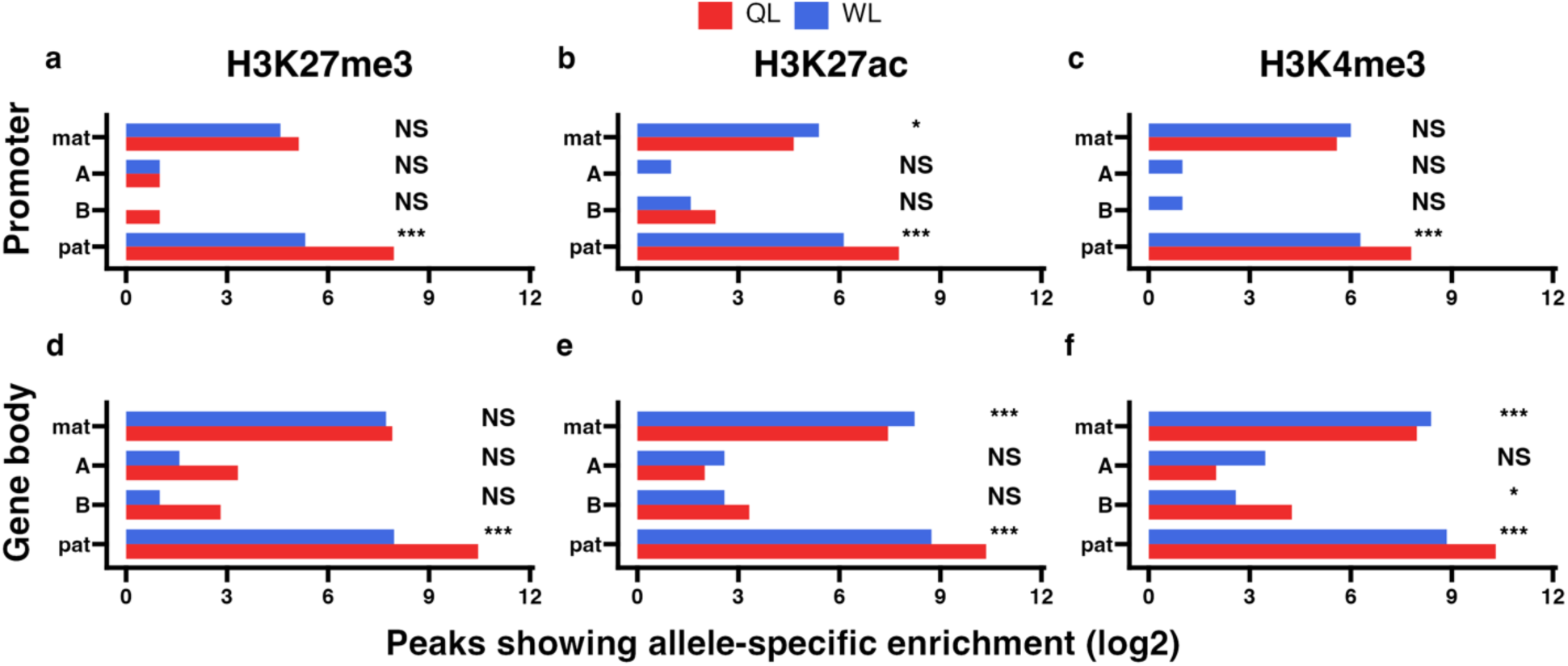
Differences in allelic enrichment of HPTMs across promoters (P) and gene bodies (GB) between queen-destined larvae (QL) and worker-destined larvae (WL) at 192hpf in Block 1. The log2 count of peaks showing significant maternal (mat), lineage A, lineage B, or paternal (pat) allele-specific enrichment across **(a-c)** promoters and **(d-f)** gene bodies at all tested SNP positions. In each panel, *p*-values for Chi-squared tests of independence for comparisons between the castes are indicated (NS = not significant, \**p*<0.01, \*\**p*<0.001, \*\*\**p*<0.001). Significance of allele-biased HPTM enrichment was determined using the overlap between two statistical tests: a general linear mixed model (GLIMMIX), and a Storer-Kim binomial exact test along with previously established thresholds of *p1*<0.4 and *p2*>0.6 for maternal bias, *p1*>0.6 and *p2*<0.4 for paternal bias, *p1*<0.4 and *p2*<0.4 for lineage B bias and *p1*>0.6 and *p2*>0.6 for lineage A bias (Wang & Clark, 2014).

### POEs on transcription are associated with POEs on HPTM enrichment

We next examined the relationships between genes with POEs on transcription and POEs on HPTM enrichment, separately for gene bodies and promoters. For each caste and Block, genes with POEs on transcription and HPTM peaks with POEs on enrichment significantly overlapped in all cases except promoter H3K27me3 in Block 2 WL (Supplementary Materials: S7.1-7.2). For all three HPTMs, across both gene bodies and promoters, for each caste and Block, POEs on HPTM enrichment were predictive of POEs on transcription, with stronger effects observed for promoters than for gene bodies (Figure 6, Supplementary Materials: S7.3). Thus, POEs on transcription co-occur with POEs on enrichment of H3K27me3, H3K4me3, and/or H3K27ac.

**Figure 6.**
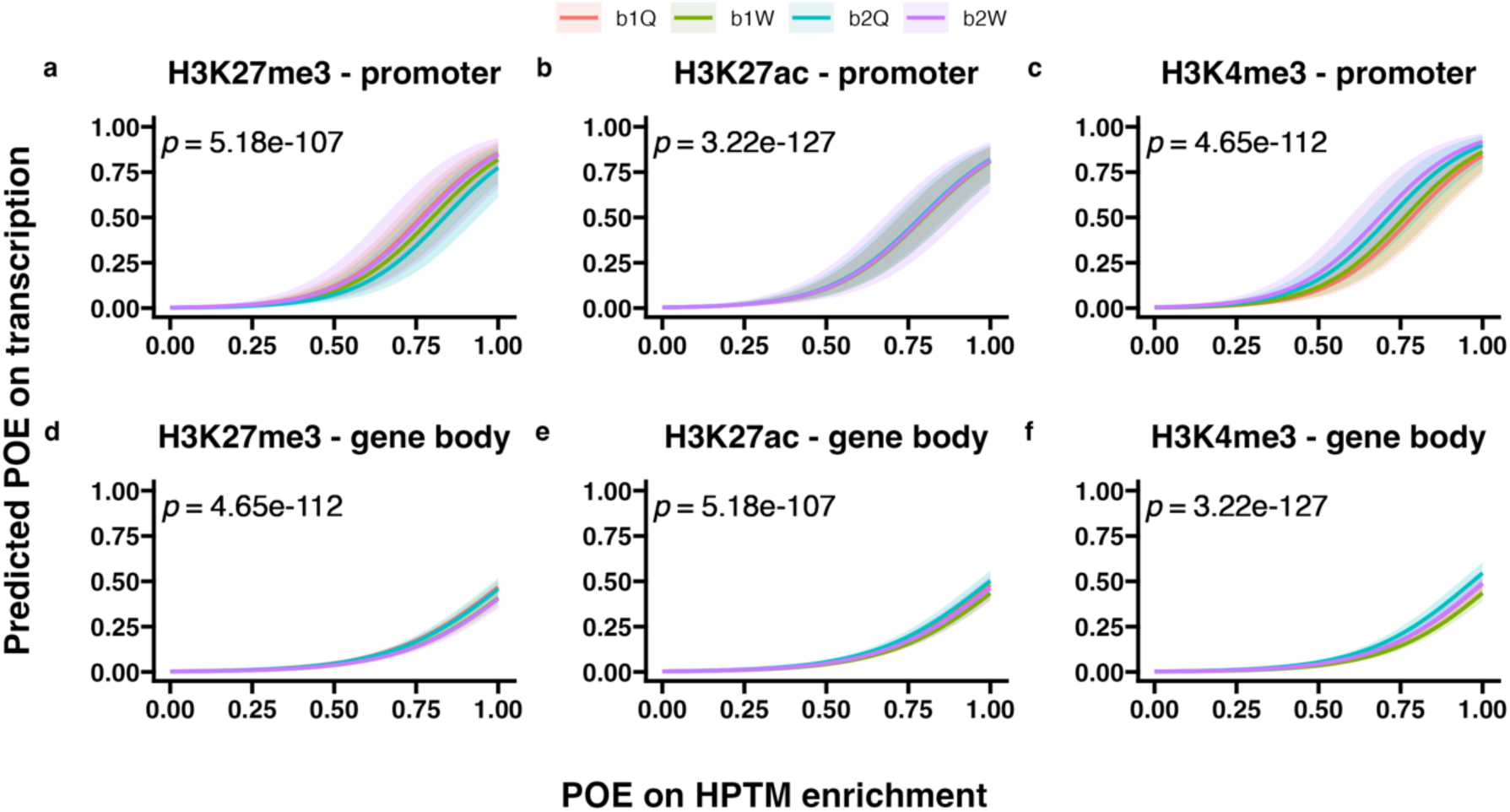
Predicted POEs on transcription given POEs on enrichment of each HPTM across promoters **(a-c)** and gene bodies **(d-f)** in each caste and Block. 95% confidence intervals are indicated. For the independent variable, genes were scored as “1” if they showed POEs on transcription or “0” if they were unbiased. For the response variable, genes were scored as “1” if a peak showing POEs on HPTM enrichment overlapped the gene or “0” if all peaks were unbiased. Results are displayed for each Block (b1 & b2) and caste (Q & W), separately. See Supplementary Materials: S7.3 for associated odds ratios tests.

Some genes showing POEs on transcription were marked by H3K27me3 on the allele with lower transcript abundance and by H3K27ac and/or H3K4me3 on the allele with higher transcript abundance, while other genes showed more complex patterns (Figure 7; Supplementary Materials: S7). For example, in QL, the *cubitus interruptus* (*Ci*) gene showed the expected patterns of allele-biased transcription and HPTM enrichment (Figure 7a), with patrigene-biased transcription, H3K27ac and H3K4me3, and matrigene-biased H3K27me3. In contrast, *frizzled 2* (*fz2*) showed patrigene-biased transcription and enrichment of all three HPTMs (Figure 7b). In WL, the *disks large 1 tumor suppressor protein* (*Dlg*) gene showed patrigene-biased transcription and matrigene-biased H3K27me3, but also peaks of both patrigene- and matrigene-biased H3K27ac and H3K4me3 (Figure 7c). Additionally in WL, *ecdysone receptor* (*Ecr*) showed patrigene-biased transcription but matrigene-biased enrichment of all three HPTMs (Figure 7d). Note that in each case, the introns of these genes contain promoters for antisense noncoding RNA (ncRNA) transcripts: *Ci* overlaps with *transfer RNA phenylalanine* (LOC107964365), *fz2* overlaps with the ncRNA *LOC102656808*, *Dlg* overlaps with multiple ncRNAs including *LOC102655405*, *LOC107965307*, and *LOC107965308*, and *Ecr* overlaps with the ncRNA *LOC107964817*.

**Figure 7.**
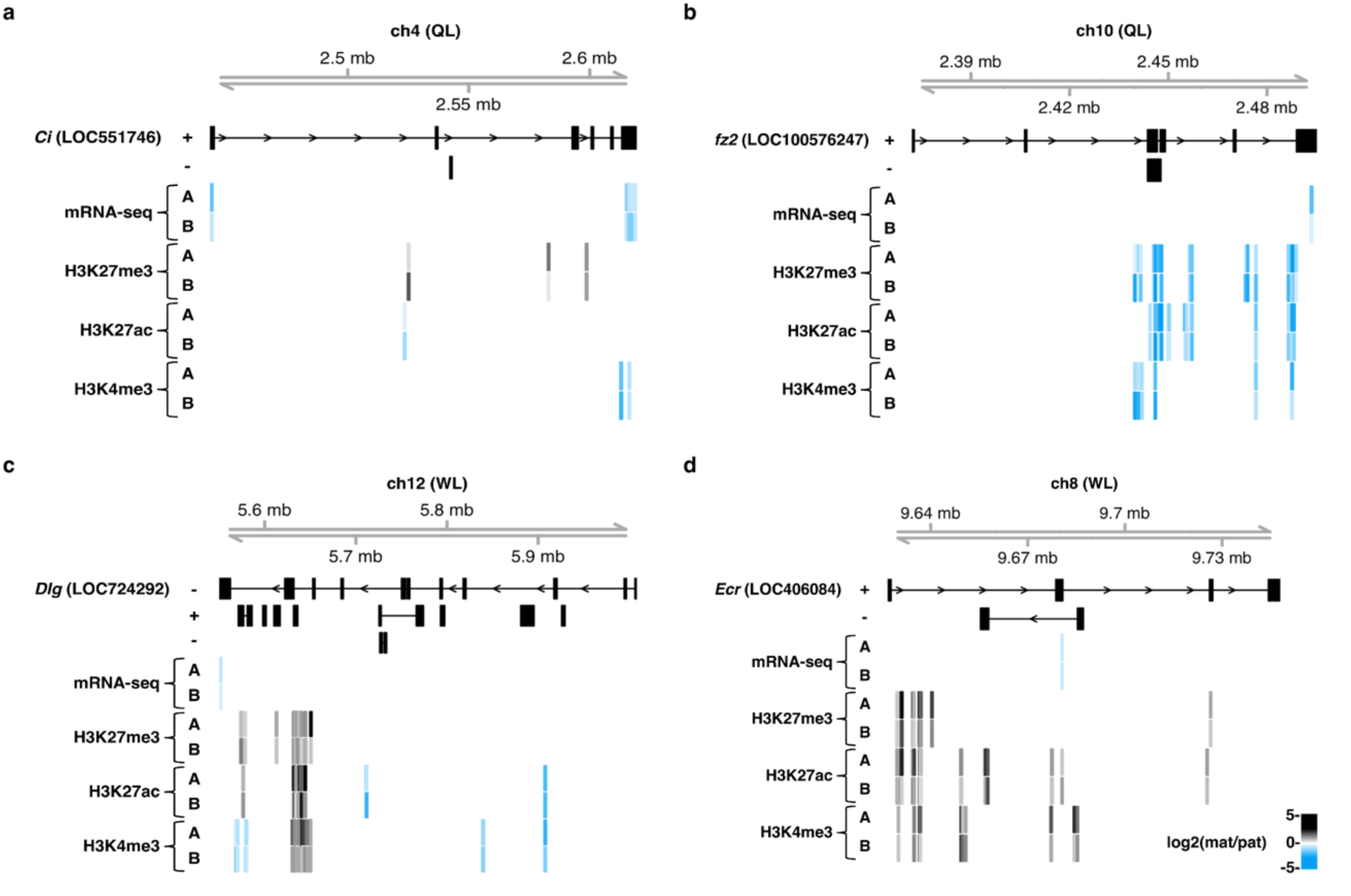
Examples of POEs on transcription and HPTM enrichment. Gene models and genomic coordinates for **(a)** *Ci*, **(b)** *fz2*, **(c)** *Dlg*, and **(d)** *Ecr*, and for overlapping noncoding RNA transcripts are displayed. Strand of each transcript relative to 5’-3’ are indicated. Colored bars indicate the matrigene (mat) to patrigene (pat) transcript abundance and HPTM enrichment ratios for QL (**a-b**) and WL (**c-d**) samples in Block 1 of each lineage (A and B) for 50bp windows containing parental SNPs used to detect POEs. Positive values indicate matrigene bias (black), and negative values indicate patrigene bias (blue).

### POEs on transcription vary throughout early honey bee development

To determine whether the POEs on transcription observed in caste determination are maintained from an earlier developmental state, we assessed POEs on transcription in 24hpf eggs in an additional set of crosses, Blocks 3 and 4 (See Methods). At approximately 60× genome coverage, we detected 364,215 (±45,418) high-confidence homozygous SNPs per F1 diploid queen and 772,598 (±39,432) per F1 haploid drone. Of these SNPs, 45,731 (±346.5) were within transcripts and varied in their sequence identity between the parents of each cross, allowing for the identification of parent-of-origin reads in the F2 eggs. The number of SNPs and transcripts examined in this study after filtering SNP positions with low mRNA-seq read coverage in the F2 eggs are listed in Supplementary Materials: S3.1-3.2.

We identified transcripts exhibiting both matrigene- and patrigene-biased expression, with more matrigene- than patrigene-biased transcripts in both Blocks (see Figure 8 for results from Block 3 and Supplementary Materials: S3.4 for results from Block 4). Overall, genes with POEs in 24hpf eggs were enriched for signaling peptides, including seven genes belonging to the *major royal jelly protein* gene family (Supplementary Materials: S8.1). Although only a small subset of genes (*n*=42) showed POEs in both developmental stages, out of the total in 192hpf larvae (*n*=508) from Blocks 1 and 2, and in 24hpf eggs (*n*=438) from Blocks 3 and 4, this overlap was statistically significant (*X^2^*-test, *p*=1×10^-5^) even though the Blocks were derived from different parents and genetic stocks.

**Figure 8.**
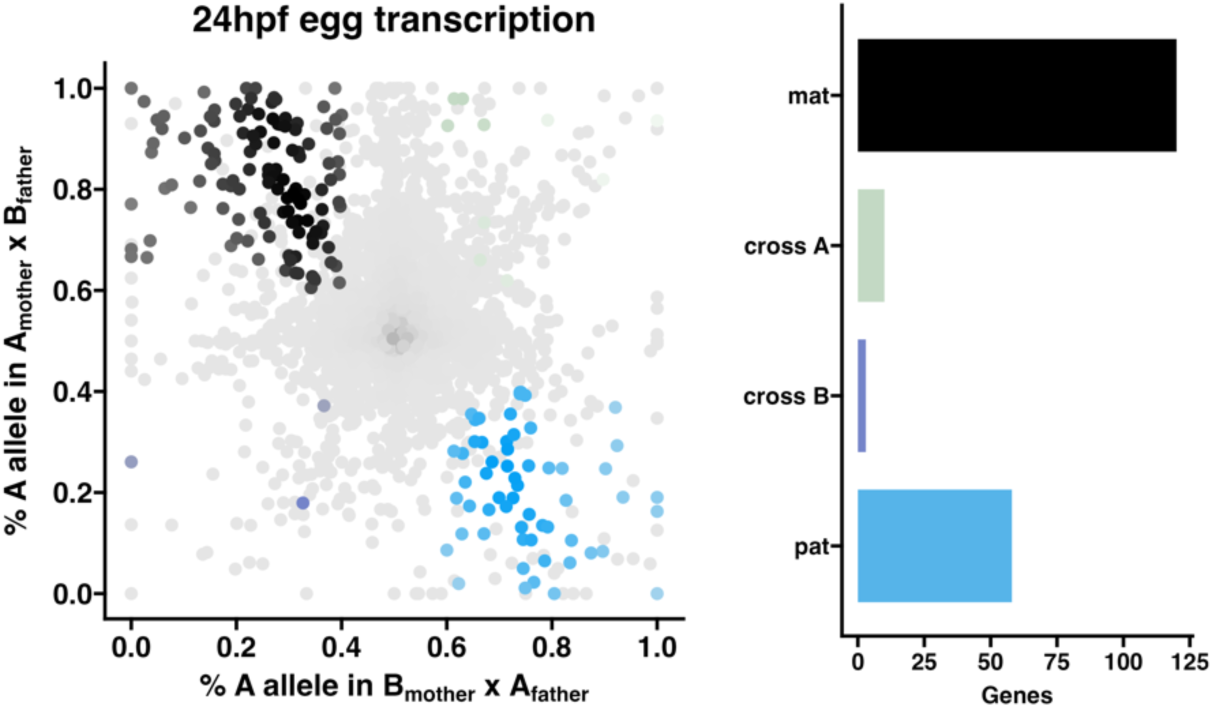
POEs on transcription in F2 eggs at 24hpf. Allele-specific transcriptomes were assessed in reciprocal crosses between different stocks of European honey bees (See Methods: Figure 10, Block 3). The x-axis represents, for each transcript, the proportion of lineage A reads in bees with a lineage B mother and lineage A father (*p1*). The y-axis represents, for each transcript, the proportion of lineage A reads in bees with a lineage A mother and lineage B father (*p2*). Each color represents a transcript which is significantly biased at all tested SNP positions: black is maternal (mat), green is lineage A, gold is lineage B, blue is paternal (pat), and grey is not significant. Bar plot (right): the number of transcripts showing each category of allelic bias. Significance of allele-biased transcription was determined using the overlap between two statistical tests: a general linear mixed model (GLIMMIX), and a Storer-Kim binomial exact test along with previously established thresholds of *p1*<0.4 and *p2*>0.6 for maternal bias, *p1*>0.6 and *p2*<0.4 for paternal bias, *p1*<0.4 and *p2*<0.4 for lineage B bias and *p1*>0.6 and *p2*>0.6 for lineage A bias (Wang & Clark, 2014). Similar results were obtained in Block 4, see Supplementary Materials: S3.3 for details.

### Genes showing parent-of-origin intragenomic conflicts in honey bee eggs and larvae have traits associated with imprinted genes in other taxa

In addition to identifying a potential gene regulatory modality associated with POEs on transcription in honey bees, we found that genes with POEs in honey bees share additional features of imprinted genes in other taxa (Figure 9) (Sandovici *et al*., 2006; Hutter *et al*., 2010; Wolf, 2013). Patrigene- or matrigene-biased transcription was consistent for a set of genes between developmental stages and castes (Figure 9a), suggesting stability of gene regulation after cell replication (Sanli & Feil, 2015). Compared to randomly sampled same-sized lists, the *n*=885 genes with POEs identified in 24hpf eggs and 192hpf larvae cluster in the genome (random clustering=28.06%, SD=±2.06%, clustering in list=46.42%, *p*=1×10^-10^; Supplementary Materials: S9.1). Compared to other genes that were expressed in these tissues and could be assessed for POEs in our study, genes with POEs are experiencing significantly higher recombination rates within *Apis* (*t*-test, *p*=0.002; Figure 9b), have longer introns (*t*-test, *p*=4.27×10^-11^; Figure 9c) with higher GC content (*t*-test, *p*=1.29×10^-12^; Figure 9d) and CpG density (*t*-test, *p*=4.04×10^-10^; Figure 9e) (Supplementary Materials: S9.2).

**Figure 9.**
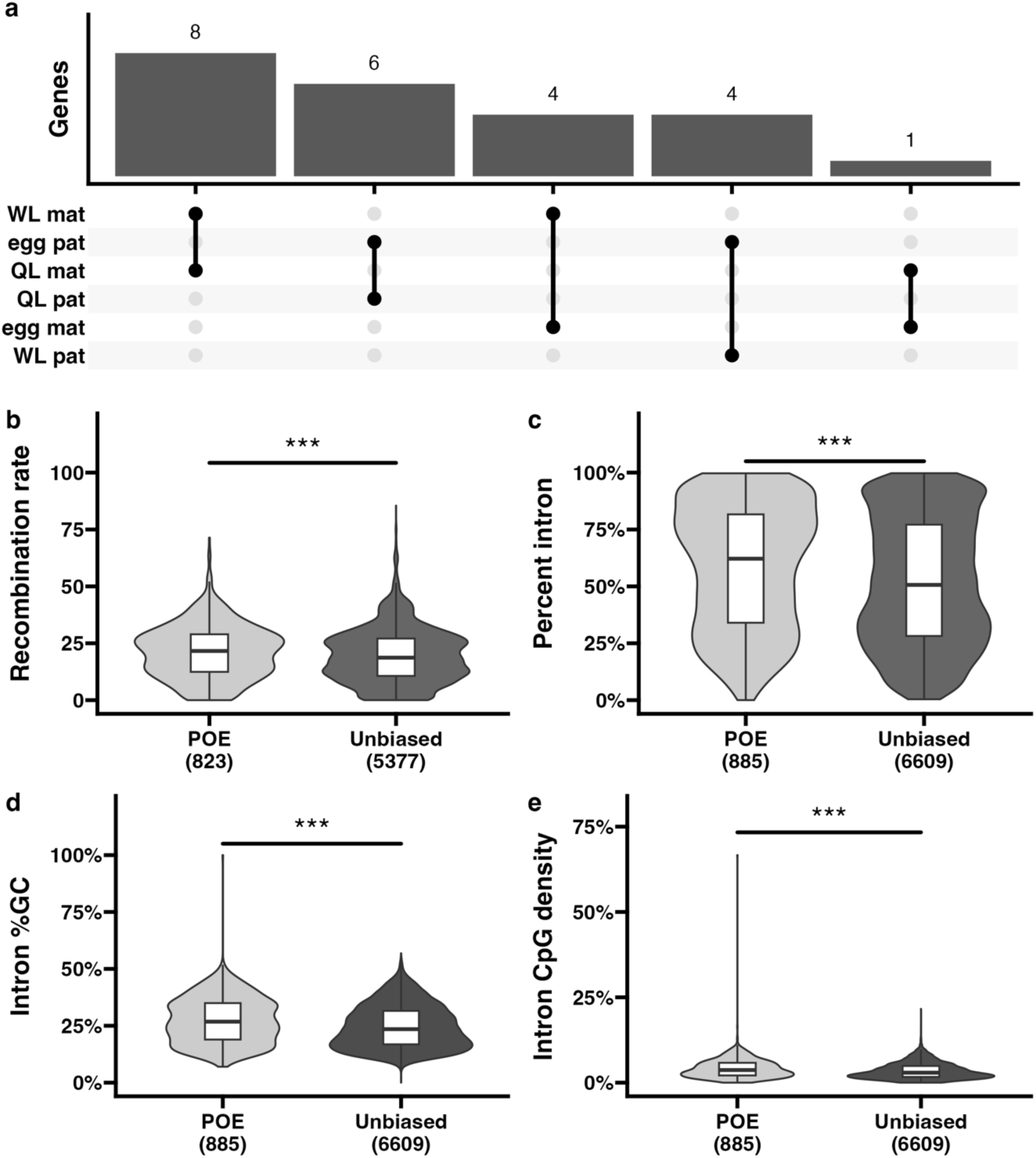
Genes with POEs on transcription in honey bees share features with imprinted genes in other taxa. **(a)** Upset plot of genes showing paternal (pat) and maternal (mat) allele-biased transcription in 24hpf eggs and 192hpf worker- and queen-destined larvae (WL and QL, respectively). **(b-e)** Comparison of **(b)** intron length, **(c)** recombination rates, **(d)** GC content, and **(e)** CpG densities between genes with POEs on transcription and unbiased genes. Significance of independent *t*-tests are indicated (\*\*\**p*<0.0001). See Supplementary Materials: S9 for details.

## Discussion

Our study provides comprehensive evidence that parent-of-origin effects (POEs) on transcription and histone post-translational modifications (HPTMs) are associated with caste determination in the honey bee *Apis mellifera*, potentially via a chromatin-regulation-based gene regulatory mechanism which has similar effects on genes (recombination rates, intron length, intron GC content and CpG density) as genomic imprinting via DNA methylation (DNAme). Across two distinct genetic Blocks, we identified hundreds of genes with POEs on transcription in both queen-destined larvae (QL) and worker-destined larvae (WL) at 192 hours post-fertilization (hpf), a developmental stage wherein the caste fates have canalized and are irreversible (Wojciechowski *et al*., 2018). Patrigene-biased transcripts were enriched in QL compared to WL, consistent with predictions of the Kinship Theory of Intragenomic Conflict (KTIC) that patrigenes should favor the reproductive (queen) caste fate to increase their transmission (Queller, 2003; Gardner & Úbeda, 2017). Genes with POEs were enriched in developmental signaling pathways and gene networks related to transcriptional regulation. A subset of genes with POEs were differentially expressed between castes and enriched for genes known to play roles in reproductive caste determination, such as *vitellogenin* and *major royal jelly protein* genes. This suggests that parent-of-origin intragenomic conflicts may have cascading effects on gene networks and directly influence dosage of key developmental regulators underpinning reproductive caste determination.

While previous theoretical work has suggested that intragenomic conflict may influence caste determination in eusocial insects (Queller, 2003; Drewell *et al*., 2011; Matsuura, 2019; Ferriera *et al*., 2024), we provide the first empirical support of this hypothesis. Patrigenes expressed in eggs (this study and Smith *et al*., 2020) may prime larvae to take advantage of worker efforts to rear new queens, perhaps hastening their development, since the first queen to emerge will attempt to kill all the other developing queens she detects within the colony (reviewed in Jackson & Robinson, 2018), or otherwise enhancing their reproductive potential as adults, such as by increasing ovariole number. Moreover, if a larva develops into a worker, patrigenes expressed in earlier stages may contribute to ovariole development (Hartfelder *et al*., 2018), resulting in an adult bee with larger ovaries (Galbraith *et al*., 2016 & 2021) that is less responsive to the “altruism”-promoting queen mandibular pheromone (Bresnahan *et al*., 2023b) and more likely to successfully compete with her sisters for ovary activation if the queen is lost (Makert *et al*., 2006).

Previous studies in social insects examining POEs on DNAme found no association with POEs on transcription (Wu *et al*., 2020; Marshall *et al*., 2020; Olney *et al*., 2021), even though this is the canonical mechanism of genomic imprinting in placental mammals, angiosperms, and other insects (MacDonald, 2012; Batista & Köhler, 2020; Hanna & Kelsey, 2021). Here, for the first time in a social insect, we found that POEs on transcription were associated with POEs on the distribution of three histone post-translational modifications (HPTMs) with functions in the regulation of transcription and *cis*-elements: H3K27me3, H3K27ac, and H3K4me3 (Bannister & Kouzarides, 2011). Genes with POEs on transcription significantly overlapped peaks with POEs on HPTM enrichment, and POEs on HPTM enrichment in promoters were highly predictive of POEs on transcription. Additionally, POEs varied between the castes. QL had more patrigene-biased enrichment of all three HPTMs and matrigene-biased H3K27me3 compared to WL, whereas WL had more matrigene-biased enrichment of H3K4me3 and H3K27ac compared to QL. This finding suggests that intragenomic conflict-regulated developmental canalization of reproductive caste fate may be associated with chromatin modifiers with roles in gene regulation and cellular differentiation (Chen & Dent, 2014). In further support of this mechanism, our gene network analysis revealed a caste determination associated module enriched for genes involved in chromatin regulation that showed POEs on transcription, which may further reenforce the POEs on these genes.

The associations between parental allele-biased enrichment of HPTMs and transcription demonstrated in our study resemble similar mechanisms of allele-specific gene regulation in *Drosophila* (Floc’hlay *et al*., 2021), and so-called “non-canonical” genomic imprinting in placental mammals and angiosperms (Batista & Köhler, 2020; Hanna & Kelsey, 2021; Raas *et al*., 2021; Soliman & Coughlan, 2024). As DNAme is not associated with transcriptional POEs in social insects (Wu *et al*., 2020; Marshall *et al*., 2020; Olney *et al*., 2021), our study provides support for the hypothesis that DNAme-independent genomic imprinting-like mechanisms should be widespread among taxa (Inoue, 2023). However, our study was limited in that we did not conduct any experiments to demonstrate a functional relationship between allele-biased enrichment of HPTMs and allele-biased transcription. Importantly, the timing of HPTM establishment, and mechanisms of HPTM maintenance across cell divisions, remains to be determined. Evidence in other taxa suggests that incomplete meiotic reprogramming of HPTMs can occur (Xia & Xie, 2020; Fukushima *et al*., 2023). Future studies in honey bees should assess HPTM profiles across developmental stages and characterize the dynamics of their establishment, maintenance and reprogramming during embryogenesis, in addition to identifying and manipulating the relevant histone modifier and remodeler proteins involved.

In mammalian and plant models of “non-canonical” DNAme-independent genomic imprinting, H3K27me3-mediated parental allele-specific transcriptional regulation via Polycomb Repressive Complex 2 (PRC2) appears to have evolved convergently (Raas *et al*., 2022; Inoue, 2023; Soliman & Coughlan, 2024). Across insects, PRC2-mediated H3K27me3 appears to play a general role in regulating hormone biosynthesis and developmental timing (Lu *et al*., 2013; Yang *et al*., 2021). Little is known about the functions of PRCs in honey bees, however, previous studies suggest that H3K27me3 is mediated by Enhancer of zeste homolog 2 (EZH2), the functional enzymatic component of PRC2 (Blackledge & Klose, 2021). In newly emerged worker bees, reduction in global levels of H3K27me3 via treatment with 3-Deazaneplanocin A (DZNep), an *S*-adenosyl-L homocysteine hydrolase inhibitor that depletes E(z) (Tan *et al*., 2007), has been shown to disrupt Notch signaling and promote ovary activity (Duncan *et al*., 2016, 2020). We found that honey bee larvae show caste-specific profiles of H3K27me3, and QL show enriched matrigene- and patrigene-biased H3K27me3 compared to WL. Some genes, such as *Ci* (Figure 7a), showing POEs were marked by H3K27me3 on the allele with lower transcript abundance and by H3K27ac and/or H3K4me3 on the opposite allele. Other genes showed more complex patterns of allelic transcription and HPTM enrichment that are difficult to interpret within the scope of this experiment. These complex patterns may be because we used whole larvae, and different cell types likely have different HPTM profiles. Alternatively, the introns of many imprinted genes in other taxa contain promoters for antisense noncoding RNA (ncRNA) transcripts (Kanduri, 2016). These antisense ncRNAs contribute to imprinted gene regulation by functioning as enhancers and through occlusion of the imprinted gene transcript. In our study, many genes showing POEs on transcription contained peaks of HPTM enrichment across antisense ncRNAs (Figure 7). A third explanation for these complex patterns is that in various human and mouse cell cultures, several developmentally repressed imprinted genes exhibit H3K27me3 and H3K4me3 bivalency on the allele that is normally expressed (McEwen & Ferguson-Smith, 2010; Young *et al*., 2011). Such bivalent marks may indicate a plastic regulatory state, such that the matrigene or patrigene is primed for rapid changes to transcriptional activity (Kumar *et al*., 2021). Future studies in honey bees would benefit from examining individual tissues or single cells, functional characterization of these antisense ncRNAs, or more fine-scaled temporal analyses.

Genes that are imprinted in other taxa via DNAme show unique sequence properties (Sandovici *et al*., 2006; Hutter *et al*., 2010). In these taxa, intron length, GC content, and recombination rates are generally correlated, with shorter introns being GC-rich (Roy *et al*., 2008; Zhu *et al*., 2009) and located in regions of high recombination (Comeron & Kreitman, 2000; Gazave *et al*., 2007; however, see Prachumwat *et al*., 2004). Imprinted genes present exceptions to these trends: they have relatively long, GC-rich introns and exhibit high rates of recombination. The differentially methylated regions (DMRs) associated with canonically imprinted genes occur at CpG sites, particularly within introns (Hutter *et al*., 2006; Wight & Werner, 2013; Martinez *et al*., 2016). Selection on intron length may generate new regulatory sequences (Tang *et al*., 2006) for controlling allelic dosage. Genomic clustering of imprinted genes facilitates their coordinated regulation (Wolf, 2013), and regulatory marks associated with imprinted regions – i.e., DNAme and H3K4me3 – can directly affect recombination (Zelkowski *et al*., 2019). Similarly, in honey bees, we found that genes with POEs on transcription have relatively long, CpG-rich introns that cluster throughout the genome and exhibit relatively high recombination rates compared to unbiased genes. Interestingly, previous studies in honey bees found no association between intron length and GC content, but that CpG-rich genes are hypomethylated (Zeng & Yi, 2010), differentially expressed between reproductive castes (Elango *et al*., 2009), and elevated recombination rates (Kent *et al*., 2012).

Additionally, the parent-of-origin transcription of many imprinted genes is persistent across cell divisions (Sanli & Feil, 2015). A subset of genes in our study maintained POEs on transcription from the egg to larval stages, potentially contributing to the development of traits that would be exhibited by adult bees. That most genes do not show persistent POEs could reflect developmental dynamics whereby parental conflicts arise and are resolved at different stages (Babek *et al*., 2015; van Ekelenburg *et al*., 2023). Alternatively, differences between Blocks regarding genetic lineages of the F0 stocks, whole genome sequencing coverage and thus detection of high-confidence homozygous SNPs could all contribute to the observed variation in POEs between developmental stages. Future studies assessing the temporal regulation of POEs should use consistent genetic backgrounds for all samples to overcome these limitations.

## Conclusions

Overall, our study demonstrates that predictions of the Kinship Theory of Intragenomic Conflict are supported during reproductive caste determination, and parent-of-origin variation in transcription underlies the extreme phenotypic plasticity associated with reproductive caste determination in social insects. Moreover, this study provides the first evidence that Polycomb repression mechanisms may mediate parent-of-origin intragenomic conflict in eusocial insects and demonstrates that this non-canonical mechanism is found across a broad range of taxa. Interestingly, though chromatin-mediated silencing is a distinct mechanism from DNAme-mediated silencing, these genes showed similar phenotypes in terms of gene structure and nucleotide content. Identifying specific cis-regulatory elements and chromatin remodeling enzymes involved could reveal if this represents genomic imprinting or a different gene regulatory process. Experimental perturbations are needed to examine the functional impacts of POEs on developmental plasticity and social behavior in honey bee colonies. Future research examining the tissue and developmental stage plasticity of these processes would help to clarify how these marks are maintained or removed to mediate phenotypic variation. Importantly, comparative studies across other eusocial lineages are needed to determine the generality of these findings.

## Methods

### Biological samples

Reciprocally crossed colonies were generated from F1 male drones and female queens derived from different founder-generation (F0) stocks of European honey bees (*Apis mellifera*) to collect F2 192-hours post-fertilization (hpf) queen- and worker-destined larvae (QL and WL, respectively) and 24hpf eggs. In studies using QL and WL, F1 reproductive females (queens) and males (drones) were sourced from two colonies of *A. m. ligustica* ‘Cordovan’ stock (colonies 1 and 2, or C1 and C2; Koehnen & Sons, Inc., Glenn, CA), one colony of mixed *A. m. ligustica* stock (C3; BeeWeaver Honey Farm, Navasota, TX), and one colony of *A. m. caucasia* stock (C4; Old Sol Apiaries). In studies using eggs, F1 queens and drones were sourced from two colonies of mixed *A. m. ligustica* stocks (C5 and C6; managed at a Penn State University apiary in University Park, PA), one colony of *A. m. caucasia* stock (C7; Old Sol Apiaries, Rogue River, Oregon), and one colony of *A. m. carnica* stock (C8; Susan Cobey, Coupeville, WA). F0 colonies were managed at a Penn State University apiary in University Park, PA, and separated into four Blocks, with two colonies from different genetic backgrounds assigned to each Block (Figure 10).

**Figure 10.**
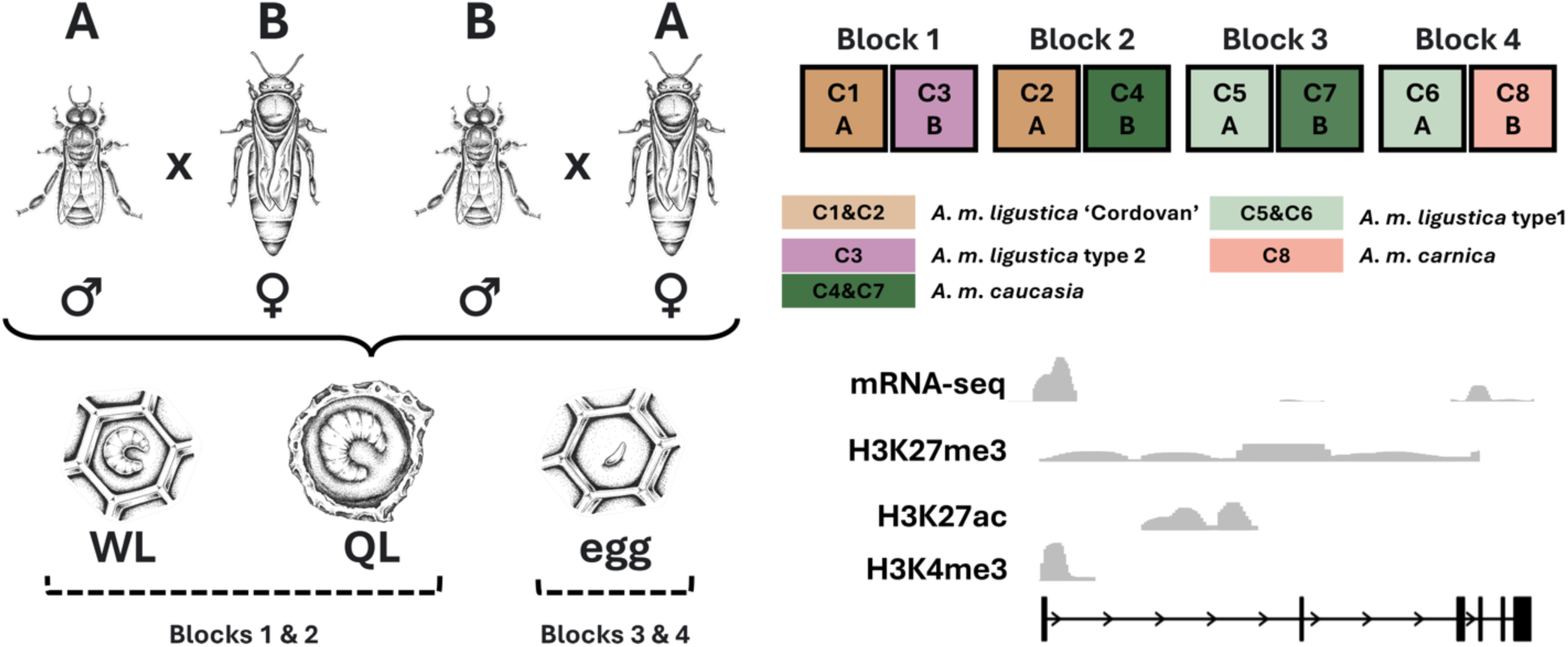
Reciprocal cross design. From each source colony (C1–C8), F1 queens (♀) and F1 drones (♂) were selected and crossed (AxB, BxA) to construct reciprocal crosses, from which F2 samples were collected. Colors indicate bee stock. Blocks 1 & 2 were generated for studies using 192-hours post-fertilization (hpf) worker-destined larvae (WL) and queen-destined larvae (QL). Blocks 3 & 4 were generated for studies using 24hpf eggs. F1 queens and drones were collected for whole genome sequencing, and F2 larvae and eggs were collected for mRNA-seq. Additionally, F2 larvae were collected for ChIP-seq of H3K27me3, H3K27ac, and H3K4me3.

From each F0 colony, F1 queens were generated using a standard commercial practice known as grafting (Connor, 2009, Anton & Grozinger, 2024). Here, a young female larva (72-96hpf) is collected with a grafting tool from a labeled brood frame confined to an emergence box with mesh cages inside the colony, on which an F0 queen was restricted to lay her eggs. The larva is placed on 1-2ul of royal jelly at the bottom of a plastic artificial queen cell attached to the cross-bar of a hive frame. Frames of grafts are then placed into a colony of worker bees from which the queen was previously removed to induce queen-rearing behavior, indicated by the building out the queen cell with beeswax and deposition of large amounts of royal jelly by the worker bees. Once the grafted larva begins to pupate and worker bees cap the queen cell with wax, the cells are moved to a standing incubator kept at a constant temperature of 33.5°C and 50% relative humidity until emergence. Adult F1 virgin queens were then housed in cages within a queenbank colony until the F1 drones reached sexual maturity. As drones tend to drift between colonies (Currie & Jay, 1991), frames of emerging F1 drone brood were identified from each F0 colony and isolated inside the colony to prevent escape. Newly emerged F1 drones were then marked on the thorax with nontoxic paint to identify their colony of origin and released back into the colony. One F1 queen (approx. 10 days old) and one F1 drone (approx. 19 days old) were selected from each colony in a Block and crossed with one F1 queen and one F1 drone from the reciprocal colony by instrumental insemination (Cobey *et al*., 2013) to create one reciprocal cross per Block (Figure 10). Instrumentally inseminated queens were labelled with a numbered tag affixed to the thorax for identification.

Each inseminated F1 queen was placed in her own mini colony of approximately 3,000 worker bees to initiate egg-laying, and the entrances of these colonies were restricted with queen excluder material to prevent escape or further mating. Approximately three weeks after the F1 queens began laying fertilized eggs, each colony was supplied with a labeled frame of empty drawn comb for the queen to lay. The following day, a transparent acrylic sheet was overlaid on each labeled frame to mark the position of newly laid eggs. From Blocks 1 and 2, upon hatching (approximately 96hpf) a subset of larvae was grafted to induce queen caste fate. After 96hrs of larval growth (approximately 192hpf), QL and WL were placed in labelled 1.5mL cyroresistant tubes, immediately frozen on dry ice, and stored at −80°C. From Blocks 3 and 4, pools of *n*=40 24hpf F2 diploid eggs were collected from labeled frames using a grafting tool, immediately dissolved in 1.5mL cyroresistant tubes containing 1mL TRIzol^®^ (Invitrogen: Waltham, MA), and stored at −80°C.

### DNA isolation and WGS library preparation

The F1 queen and drone parents of each reciprocal cross from *n*=4 Blocks (*n*=16 samples in total) were used for whole genome sequencing (WGS). The thorax of each bee was removed, and the flight muscle tissue was dissected on a platform surrounded by dry ice to keep the tissue frozen until the DNA isolation procedure. Genomic DNA was isolated using a DNeasy^®^ Blood & Tissue Kit (Qiagen: Germantown, MD). Samples were sent to the Penn State Genomics Core Facility for library preparation using NEBNext^®^ Ultra^TM^ II DNA Library Prep kits for Illumina^®^ with NEBNext^®^ Multiplex Oligos (New England Biolabs, NEB: Ipswich, MA). Each library was prepared from 500ng DNA, PCR amplified for 3 cycles, purified with AMPure XP Beads (Beckman Coulter: Brea, CA), and pooled at equimolar concentration. Pooled libraries with unique 5′ and 3′ index adapter pairs were sequenced on an NextSeq 2000 P3 flow cell (Illumina^®^: San Diego, CA) to generate 66M (±12M) 150-bp paired-end reads per library (Supplementary Materials: S1.1).

### RNA isolation and mRNA-seq library preparation

Five each of 192hpf F2 WL and QL per cross per Blocks 1 and 2 were selected for mRNA-seq (*n*=40 samples total). Whole larvae were dissolved in 1.5mL microcentrifuge tubes containing 500ul TRIzol^®^. Total RNA was isolated via chloroform extraction followed by precipitation with isopropanol and treated with 1ul (2U/ul) TURBO^TM^ DNase (Invitrogen). We assessed RNA integrity by RNA ScreenTape Analysis on a TapeStation 4150 (Agilent: Santa Clara, CA), requiring an RNA integrity number equivalent (RIN^e^) of at least 9.0/10.0 for library preparation. In total, we prepared *n*=40 sequencing libraries from 1µg total RNA per sample using Illumina^®^ Stranded mRNA Prep kits with unique 5′ and 3′ index adapter pairs (IDT for Illumina^®^ RNA UD Indexes, Set A). Libraries were PCR amplified for 10 cycles, then cleaned to remove reagents, enzymes and excess adapters using AMPure XP Beads on a magnetic separation rack (Sergi Lab Supplies: Seattle, WA). Next, libraries were assessed by D1000 DNA ScreenTape Analysis on a TapeStation 4150 to estimate concentration and determine average insert length, diluted to approximately 2nM, and quantified using a NEBNext^®^ Library Quant Kit for Illumina^®^ (NEB) on a QuantStudio^TM^ 5 Real-Time PCR System (ThermoFisher: Waltham, MA). Finally, libraries were pooled at equimolar concentration, treated with Illumina^®^ Free Adapter Blocking Reagent, cleaned with AMPure XP Beads and quantified by qPCR as before. We performed 50-bp paired-end sequencing through the Penn State Genomics Research Incubator on a NextSeq 2000 P3 flow cell to generate 56M (±14M) reads per library (Supplementary Materials: S1.2).

Three pools of *n*=40 24hpf F2 eggs per cross per Blocks 3 and 4 (*n*=12 pools in total) were selected for mRNA sequencing (mRNA-seq). Total RNA was isolated as before. Samples were sent to the Penn State Genomics Core Facility for library preparation using Illumina^®^ Stranded mRNA Prep kits with IDT for Illumina^®^ RNA UD Indexes. Each library was prepared from 500ng RNA, PCR amplified for 10 cycles, and pooled at equimolar concentration. Pooled libraries with unique 5′ and 3′ index adapter pairs were sequenced on a NextSeq 2000 P3 flow cell to generate approximately 21M (±4.5M) 150-bp single-end reads per library (Supplementary Materials: S1.2).

### Chromatin preparation

Chromatin was extracted from pools of *n*=6 192hpf F2 larvae, two pools per caste (QL and WL) per cross per Blocks 1 and 2 (*n*=16 pools in total), adapted from a protocol established for honey bee larvae described in Wojciechowski *et al*. (2018). Larvae were placed in 1.5mL microcentrifuge tubes containing 1mL of 5% cOmplete^TM^ EDTA-free protease inhibitor cocktail (PIC) (Roche Diagnostics Corporation: Indianapolis, IN) in 1× Phosphate Buffered Saline (PBS) at 4°C. Cell suspensions were produced by disaggregating the thawing tissue with RNase-Free disposable pellet pestles (Fisher Scientific: Hampton, NH) and passaging 5× through sterile 18- and 21-guage needles using 1mL Luer lock syringes. Cell viability was assessed by staining 1ul of the cell suspension in a 10ul solution of 0.4% trypan blue in 1× PBS and counting the number of unstained cells on a hemocytometer slide under a compound light microscope.

From each pool of larvae, approximately 30M cells were cross-linked in 1mL solution (1% formaldehyde, 1% PIC, 1× PBS) in a tube rotator at room temperature for 8min, followed by treatment with 100ul 2.5M glycine for 5min to stop the reaction. Cross-linked cells were then pelleted at 2,000×g for 5min at 4°C, washed twice with 400ul wash buffer A (1mL cold 5% PIC in 1× PBS), and lysed in 400ul cell lysis buffer (10mM Tris-HCl pH 8.0, 10mM NaCl, 0.2% NP-40, 1% PIC) on ice for 10min. Nuclei were then pelleted at 2,000×g for 5min at 4°C, washed twice with wash buffer A, and lysed in 500ul nuclear lysis buffer (50mM Tris-HCl pH 8.0, 10mM EDTA, 1% SDS, 1% PIC) on ice for 10min. The chromatin suspension was then diluted in 1mL immunoprecipitation (IP) dilution buffer (IPDB) (20mM Tris-HCl, 2mM EDTA, 150mM NaCl, 1% Triton X-100, 1% PIC) and sheared to 100-600bp fragments (average 250bp) at 4°C in a Bioruptor^®^ Pico sonication device (Diagenode: Denville, NJ) for 5 cycles (30 sec on, 30 sec off). Fragment length was assessed by D1000 DNA ScreenTape Analysis on a TapeStation 4150 before proceeding to the IP reaction (see Supplementary Materials: S1.3 for representative examples). From each sample, 5% (75ul; ∼2µg) aliquots were combined with 2ul DNase-free RNase A (10mg/mL) and 16ul 5M NaCl for 1hr at 65°C to remove RNA contamination, followed by 3ul (20mg/mL) proteinase K for 1hr at 45°C to reverse cross-links. Aliquots were then purified using a PCR purification kit (Qiagen), eluting in 50ul room temperature buffer EB (10mM Tris-HCl pH 8.5).

### Chromatin immunoprecipitation and ChIP-seq library preparation

Chromatin was pre-cleared by centrifugation at 14,000×g for 15min at 4°C and the supernatant was moved to a 1.5mL DNA LoBind^®^ tube (Eppendorf: Hamburg, Germany). From each chromatin sample per phenotype per lineage, 5% (75ul;∼2µg) was removed and combined to a separate tube for the corresponding input (no-IP) sample. For each IP, antibody:bead complexes of 5ul (1µg/uL) antibody and 25ul (30mg/mL) Protein G Dynabeads^TM^ (Invitrogen) were prepared. To reduce non-specific binding, beads were added to 1.5mL DNA LoBind^®^ tubes, placed on a magnetic rack to remove the supernatant, and combined with 1mL cold blocking solution (1× PBS and 0.5% bovine serum albumin). Beads in blocking solution were vortexed briefly, placed back on the magnetic rack to remove the supernatant, washed with 500ul blocking solution, then combined with 50ul cold 1× PBS. Antibodies previously used successfully in ChIP studies in honey bees against H3K27me3 (Abcam #ab6002: Cambridge, UK) (Duncan *et al*., 2020), H3K27ac (Active Motif #39133: Carlsbad, CA) and H3K4me3 (Abcam #ab12209) (Wojciechowski *et al*., 2018) were then added to separate tubes containing 50ul blocked protein G beads and rotated at 4°C for 6hrs.

Antibody:bead complexes were placed on a magnetic rack to remove the supernatant, combined with 450ul (∼12µg) of pre-cleared chromatin preparation and rotated at 4°C for 6hrs. Samples were then placed back on the magnetic rack to remove unbound chromatin, washed once with 500ul cold wash buffer B (50mM Tris-HCl pH 8.0, 150mM NaCl, 0.1% SDS, 1% NP-40, 0.5% sodium deoxycholate), once with 500ul cold wash buffer C (50mM Tris-HCl pH 8.0, 250mM NaCl, 1mM EDTA, 0.1% SDS, 1% NP40, 0.5% sodium deoxycholate), once with cold wash buffer D (50mM Tris-HCl pH 8.0, 250mM LiCl, 1mM EDTA, 1% NP-40, 0.5% sodium deoxycholate) and once with cold TE buffer (0.5M EDTA, 1M Tris-HCl pH 8.0). Bound chromatin was eluted from the beads with 200ul room temperature elution buffer (1% SDS, 100mM NaHCO_3_), and then treated to remove RNA contamination and to reverse cross-links, purified using a PCR purification kit and eluted in 50ul room temperature buffer EB as before.

We prepared ChIP libraries for 2 biological replicates (pooled larvae) x 2 castes x 2 crosses x 2 Blocks x 3 histone post-translational modifications (HPTMs) = 48 libraries, plus 1 input DNA library x 2 castes 2 x crosses x 2 Blocks = 8 libraries. In total, we prepared *n*=56 ChIP-seq libraries using NEBNext^®^ Ultra^TM^ II DNA Library Prep kits for Illumina^®^ with NEBNext^®^ Multiplex Oligos. Input sample libraries were prepared from 200ng DNA and PCR amplified for 5 cycles. ChIP sample libraries were prepared from 1-10ng DNA and PCR amplified for 12 cycles. Library cleanup, fragment length analysis, dilution, quantitation, pooling, and treatment to remove free adapters was conducted following the same procedure as in mRNA library preparation. We performed 50-bp paired-end sequencing through the Penn State Genomics Research Incubator on a NextSeq 2000 P3 100-cycle flow cell to generate approximately 49M (±26M) reads library (Supplementary Materials: S1.3).

### Differential gene expression analysis

We used Kallisto (Bray *et al*., 2016) with the setting “-rf-stranded” to estimate transcript abundances of larval samples. A transcriptome index was generated with gffread (Pertea & Pertea, 2020) using the Amel_HAv3.1 RefSeq genome feature file. Transcript abundance files were combined into a gene expression matrix using the tximport package (Soneson *et al*., 2022) in R (R Core Team, 2023), with the “countsFromAbundance” argument set to “lengthScaledTMP” to adjust for transcript length. A transcript-to-gene table was constructed from the RefSeq annotation using the makeTxDbFromGFF function from the GenomicFeatures package (Lawrence *et al*., 2013). Following variance stabilizing transformation (VST) of the gene expression matrix, sample relationships were assessed using *k*-means clustering with *k*=2. We used DESeq2 (Love *et al*., 2014) and a two-factor design to test for differential expression between castes, treating lineage as a cofactor. Effect sizes were adjusted using Bayesian shrinkage estimators with the apeglm package (Zhu *et al*., 2019). For this study, we focused on differentially expressed genes at thresholds of |log_2_(fold change, FC)| >1.5 and false discovery rate (FDR) <0.01 (Supplementary Materials: S2.1).

### Generation of F1 genomes and identification of SNPs within transcripts

WGS reads were trimmed with fastp (Chen et al. 2018) to remove adapter sequences and filter low-quality reads, then aligned to the *A. mellifera* Amel_HAv3.1 genome assembly (Wallberg *et al*., 2019) with BWA-MEM (Li, 2013) using the default settings. Alignments were coordinate sorted and filtered using samtools (Danecek *et al*., 2021) to remove unmapped reads, reads with unmapped mates, secondary and supplementary alignments. Duplicate alignments were removed with GATK MarkDuplicates (McKenna *et al*., 2010). Variants were detected using freebayes (Garrison & Marth, 2012) to account for differences in ploidy between diploid queens and haploid drones. The variant calls were filtered for quality and read depth with VCFtools (Danacek *et al*., 2011) using the settings “--minQ 30 --min-meanDP 10 --min-DP 10 --max-meanDP 50 -- max-DP 50”, and all variants except homozygous SNPs were filtered with samtools BCFtools. The high-confidence homozygous SNPs were then integrated into the Amel_HAv3.1 genome assembly with GATK FastaAlternativeReferenceMaker, and intersected using bedtools (Quinlan & Hall, 2010) to identify SNPs that were unique to each parent but shared between the crosses of a reciprocal cross pair. Parent-specific SNPs were then intersected with the RefSeq *A. mellifera* gene annotation to identify positions within the longest transcript for each gene using the GenomicFeatures package in R. Transcripts for miRNA and tRNA genes were filtered, as these loci are inappropriate for allele-specific transcriptomic analysis using short sequencing reads (Wang & Clark, 2014).

### Allele-specific transcriptome analysis

mRNA-seq reads were trimmed with fastp to remove adapter sequences and filter low-quality reads, then aligned to each respective parent genome using STAR (Dobin *et al*., 2013). Alignments with >0 mismatches, unmapped mates and secondary alignments were filtered using samtools. For each sample, read coverage was calculated at each SNP using bedtools intersect in strand-aware mode. Coverage files were then combined in R to create a matrix of F2 mRNA-seq read counts at F1 SNPs within *A. mellifera* transcripts. Whenever two or more SNPs received the same counts across all samples within a transcript (as the distance between each SNP was shorter than the length of the reads which aligned to them), one SNP was chosen at random. Finally, SNPs that received <10 counts and transcripts with <2 SNPs were filtered out prior to downstream analyses. The number of SNPs and transcripts examined in this study after filtering SNP positions with low mRNA-seq read coverage in the F2 larvae, and number of transcripts showing parent-of-origin effects (POEs) on transcription in each Block are listed in Supplementary Materials: S3.1-3.2.

We utilized the Storer-Kim (SK) binomial exact test of two proportions (Storer & Kim, 1990; Wang & Clark, 2014) and a general linear mixed effects model with interaction terms (GLIMMIX) to identify transcripts that exhibited POEs on expression following methods described previously (Kocher *et al*., 2015; Galbraith *et al*., 2016; Galbraith *et al*., 2021; Bresnahan *et al*., 2023). To adjust for variation in sequencing depth between samples, we 1) fit the models to the raw read counts and specified library size factors as an offset term, and 2) used normalized read counts for the SK tests. Library size factors were estimated using the median of ratios normalization (MRN) method with the DESeq2 package on the within-sample SNP-wise sums of allelic read counts (Castel *et al*., 2015; Fan *et al*., 2021), treating lineage and phenotype as cofactors. Normalized read counts were generated by dividing the raw read counts by these size factors.

SK tests were conducted for each transcript for each SNP to test for significant differences in the proportion of maternal and paternal read counts. Test results were then FDR corrected and aggregated by transcript. For each transcript, all positions were required to exhibit the same direction of parent- or lineage-specific biases of *p1* (the proportion of reads mapping to the “A” allele in offspring from the “B” queen and “A” drone), and *p2* (the proportion of reads mapping to the “A” allele in offspring from the “A” queen and “B” drone). As most candidate genes with allele-specific expression ratios between 0.4 and 0.6 will not validate across multiple individuals (Wang & Clark, 2014), we chose thresholds of *p1*<0.4 & *p2*>0.6 for maternal bias, *p1*>0.6 & *p2*<0.4 for paternal bias, *p1*<0.4 & *p2*<0.4 for lineage B bias, and *p1*>0.6 & *p2*>0.6 for lineage A bias. Additionally, a GLIMMIX was fit for each transcript to test for an effect of parent, lineage, and their interaction, offset by the log of the library size factors, and treating SNP and individual as random effects. Wald tests were performed to test for significance of the coefficients. Test results were FDR corrected, and only transcripts with a significant effect of parent or lineage, but not their interaction, were considered to exhibit bias. To avoid identifying POE genes influenced by lineage-of-origin effects (LOEs), or LOE genes influenced by POEs, we excluded genes that showed a significant interaction effect even if this was not the main effect. Only genes considered to exhibit bias in both tests were reported as exhibiting bias. Chi-squared tests were performed to determine whether there was an enrichment for allelic bias (paternal, maternal, lineage A or B) in QL relative to WL.

### Weighted Gene Co-expression Network Analysis (WGCNA)

WGCNA of the larval transcriptomes was performed in R (Langfelder & Horvath, 2008). Unsigned co-expression network modules were constructed using the VST gene expression matrix and a soft-threshold power of 15 to balance scale independence and mean connectivity. Modules with eigengene correlation coefficients >0.75 were merged. We then calculated correlation coefficients of each module eigengene with caste (WL=0, QL=1). Hypergeometric tests were conducted to identify modules that were enriched for POE genes (Supplementary Materials: S2.2).

### ChIP-seq analysis

ChIP-seq reads were trimmed with fastp to remove adapter sequences and filter low-quality reads with settings “-l 20 -e 20”. The filtered reads were then aligned to the *A. mellifera* Amel_HAv3.1 genome assembly with bowtie2 (Langmead & Salzberg, 2012) with settings “-I 50 -X 800 --no-mixed --no-discordant”. Low-quality alignments (Q<30) were filtered with samtools, and duplicates were removed with gatk MarkDuplicates. Alignments to problematic regions (e.g., repeats and areas of high background noise) (Amemiya *et al*., 2019) were filtered using a blacklist generated by combining repetitive regions identified with RepeatMasker (Smit *et al*., 2015) and high background noise regions identified with the input libraries using GreyListChIP (Brown, 2023). Cross-correlation analyses were performed using phantompeakqualtools (Landt *et al*., 2012) to estimate predominant aligned fragment length, and to calculate the normalized strand cross-correlation coefficient (NSC) and relative strand cross-correlation coefficient (RSC), requiring NSC>1 and RSC>1 to proceed with downstream analyses. (see Supplementary Materials: S1.3).

Broad peaks were called using MACS2 (Zhang *et al*., 2008) with genome size set to 2.25×10^8^. For quality control, peak distributions around transcription start sites (TSS) and genomic features were visualized using ChIPseeker (Wang *et al*., 2022) (Supplementary Materials: S4.1). A set of consensus peaks were then called by pooling together the signal libraries for each HPTM by caste and Block to identify regions for allelic enrichment analysis. The reads were then aligned to each respective parent genome with bowtie2 to identify allelic enrichment within consensus peaks following the same procedure as with the mRNA-seq reads and transcripts. Separate analyses were conducted for promoters and gene bodies, distinguished using GenomicFeatures. Peaks were then assigned to overlapping genes using findOverlaps from the GenomicRanges package (Lawrence *et al*., 2013) in R. The number of SNPs and peaks examined in this study after filtering SNP positions with low ChIP-seq read coverage in the F2 larvae, and number of peaks showing POEs on HPTM enrichment in each Block are listed in Supplementary Materials: S5.1-5.2.

Differences in genome-wide enrichment of each HPTM were determined using csaw (Lun & Smyth, 2016) for 1kb sliding windows with a stride of 150bp across the full length of the Amel_HAv3.1 genome assembly. Fragment abundance for all signal and input libraries was calculated for each window, and following normalization to 1x genome coverage, windows where fragment abundance of the signal was log_2_(FC) <1 over the input were filtered prior to analysis. Statistical tests of differential enrichment between the castes was performed with edgeR (Robinson *et al*., 2010), treating lineage as a cofactor. Overlapping windows were then merged into regions of up to 10kb to calculate per-region fold changes and FDR corrected *p*-values, requiring thresholds of |log_2_(FC)| >0 and FDR <0.1 for differential enrichment. Results of these analyses are available in Supplementary Materials: S4.2.

Logistic regressions were conducted to assess relationships between ChIP enrichment and transcription. For the independent variable, the transcript abundance estimates were pooled across samples for each caste and Block, separately, normalized using the variance stabilizing transformation, and standardized to a range of 0<*x*<1. For the response variable, genes were scored as “1” if a peak overlapped the gene or “0” if no peak overlapped the gene. Separate analyses were conducted for each HPTM, across promoters and gene bodies, respectively. We used bedtools to generate random sets of genomic intervals of the same quantity and average length in each sample group for comparison. Additional logistic regression tests were performed to assess relationships between POEs on transcription and POEs on HPTM enrichment. For the independent variable, genes were scored as “1” if they showed a POE on transcription or “0” if they were biallelic. For the response variable, genes were scored as “1” if a peak showing POEs on HPTM enrichment overlapped the gene or “0” if all peaks overlapping the gene were biallelic. Logistic regression summary tables and odds ratio test results are available in Supplementary Materials: S6-S7.

### Gene ontology (GO) enrichment analyses

GO enrichment analyses were conducted with DAVID (Sherman *et al*., 2022). We focused on biological processes terms and KEGG pathways significantly overrepresented among candidate gene sets at FDR <0.05 (Supplementary Materials: S8).

### Analyses of sequence features

Spatial clustering analysis of genes across the *A. mellifera* genome was performed using Cluster Locator (Obregón *et al*., 2018) with a maximum gap of 1. Statistical significance of spatial clustering of genes showing POEs on transcription in eggs and larvae was determined by comparison with 1,000 random lists. Intron length, GC content and CpG density of *A. mellifera* genes was calculated using GenomicFeatures. Recombination rates within *Apis* were retrieved from Slater *et al*. (2022). Comparisons between genes with POEs on transcription and unbiased genes were performed with independent *t*-tests. Results of these analyses are available in Supplementary Materials: S9.

## Declarations

### Ethics approval and consent to participate

Not applicable.

### Consent for publication

Not applicable.

### Availability of data and materials

Supplementary materials for this study, including analysis scripts for reproducing the results presented here are available at https://github.com/sbresnahan/IGC-CD-SM. All mRNA-seq, WGS, and ChIP-seq reads generated for this study are deposited in the NCBI Sequence Read Archive under BioProject accession PRJNA1106847.

### Authors’ contributions

STB and CMG designed the study. KA identified F0 bee stocks and performed instrumental inseminations. KA and STB managed experimental colonies. STB collected biological samples, performed DNA and RNA isolation, ChIP-seq, library preparation, sequencing, and conducted all bioinformatic and statistical analyses. SM and BH consulted on the bioinformatic analyses. STB wrote the initial draft of the manuscript and all authors contributed to its revision.

## Acknowledgements

We thank Kaleena Cazajkowski (Penn State) for assistance with creating and managing the honey bee stocks used in this study. Additionally, we thank Daniela James and Cheryl Keller for training on library preparation and sequencing, and assistance with optimizing our ChIP protocol. We thank the Penn State Genomics Research Incubator for providing the opportunity, facilities, and equipment for training on library preparation and sequencing. We are grateful to the Penn State Huck Institutes of the Life Sciences Genomics Core Facility (RRID:SCR_023645) for conducting whole genome sequencing of the parental stocks. Finally, we thank members of the Grozinger laboratory for critical reading of the manuscript.

## References

Al Adhami, H., Evano, B., Le Digarcher, A., Gueydan, C., Dubois, E., Parrinello, H., Dantec, C., Bouschet, T., Varrault, A., & Journot, L. (2015). A systems-level approach to parental genomic imprinting: The imprinted gene network includes extracellular matrix genes and regulates cell cycle exit and differentiation. Genome Research, 25, 353–367.

Alhosin, M. (2023). Epigenetics mechanisms of honeybees: Secrets of royal jelly. Epigenet Insights, 16, 25168657231213717.

Amemiya, H. M, Kundaje, A., Boyle, A. P. (2019). The ENCODE Blacklist: Identification of problematic regions of the genome. Scientific Reports, 9, 9354.

Anton, K., Grozinger, C. (2024). Queen cell production: Grafting and graft-free methods. University Park, PA; Penn State Extension. 10.26207/65z4-pw38

Ashby, R., Forêt, S., Searle, I., Maleszka, R. (2016). MicroRNAs in honey bee caste determination. Scientific Reports, 6, 18794.

Beacon, T. H., Delcuve, G. P., López, C., Nardocci, G., Kovalchuk, I., van Wijnen, A. J., Davie, J. R. (2021). The dynamic broad epigenetic (H3K4me3, H3K27ac) domain as a mark of essential genes. Clinical Epigenetics, 13, 138.

Babek, T., DeVeale, B., Tsang, E. K., Zhou, Y., Li, X., Smith, K. S., et al. (2015). Genetic conflict reflected in tissue-specific maps of genomic imprinting in human and mouse. Nature Genetics, 47, 544–549.

Bannister, A. J., Kouzarides, T. (2011). Regulation of chromatin by histone modifications. Cell Research, 21, 381–395.

Barchuk, A. R., Cristino, A. S., Kucharski, R., Costa, L. F., Simões, Z. L. P., Maleszka, R. (2007). Molecular determinants of caste differentiation in the highly eusocial honeybee *Apis mellifera*. BMC Dev Biol, 7, 70.

Batista, R. A., Köhler, C. (2020). Genomic imprinting in plants—revisiting existing models. Genes & Development, 34, 24–36.

Blackledge, N. P., Klose, R. J. (2021). The molecular principles of gene regulation by Polycomb repressive complexes. Nat Rev Mol Cell Biol, 22, 815–833.

Bowman, G. D., Poirier, M. (2015). Post-translational modifications of histones that influence nucleosome dynamics. Chem Rev, 6, 2274–2295.

Bray, N. L., Pimentel, H., Melsted, P., Pachter, L. (2016). Near-optimal probabilistic RNA-seq quantification. Nat. Biotechnol, 34, 525–527.

Bresnahan, S. T., Lee, E., Clark, L., Ma, R., Rangel, J., Grozinger, C. M., Li-Byarlay, H. (2023a). Examining parent-of-origin effects on transcription and RNA methylation in mediating aggressive behavior in honey bees (*Apis mellifera*). BMC Genomics, (24), 315.

Bresnahan, S. T., Ma, R., Galbraith, D. A., Rangel, J., Grozinger, C. M. (2023b). Beyond conflict: Kinship theory of intragenomic conflict predicts individual variation in altruistic behavior. Molecular Ecology, (32), 5823–5837.

Brown, G. (2023). GreyListChIP: Grey lists – mask artefact regions based on ChIP inputs. R package version 1.34.0. Available from: https://bioconductor.org/packages/GreyListChIP.

Bogan, S. N., Yi, S. V. (2024). Potential role of DNA methylation as a driver of plastic responses to the environment across cells, organisms, and populations. Genome Biol & Evol, 16(2), evae022.

Castel, S. E., Levy-Moonshine, A., Mohammadi, P., Banks, E., Lappalainen, T. (2015). Tools and best practices for data processing in allelic expression analysis. Genome Biol, 16, 195.

Chen, T., Dent, S. Y. R. (2014). Chromatin modifiers and remodelers: regulators of cellular differentiation. Nature Reviews Genetics, 15, 93–106.

Chen, S., Zhou, Y., Chen, Y., Gu, J. (2018). fastp: an ultra-fast all-in-one FASTQ preprocessor. Bioinformatics, 34, i884–i890.

Cobey, S. W., Tarpy, D. R., Woyke, J. (2013). Standard methods for instrumental insemination of *Apis mellifera* queens. J. Apic. Res, 52, 1–18.

Comeron, J. M., Kreitman, M. (2000). The correlation between intron length and recombination in drosophila. Dynamic equilibrium between mutational and selective forces. Genetics, 156(3), 1175–90.

Connor, L. (2009). Queen rearing essentials. 1st ed. Wicwas Press, LLC.

Currie, R. W., Jay, S. C. (1991). Drifting behaviour of drone honey bees (*Apis mellifera* L.) in commercial apiaries. J. Apic. Res, 30, 61–68.

Danacek, P., Auton, A., Abecasis, G., Albers, C. A., Banks, E., et al. (2011). The variant call format and VCFtools. Bioinformatics, 27(15), 2156–2158.

Danecek, P., Bonfield, J. K., Liddle, J., Marshall, J., Ohan, V., Pollard, M. O., Whitwham, A., Keane, T., McCarthy, S. A., Davies, R. M., et al. (2021). Twelve years of SAMtools and BCFtools. GigaScience, 10, giab008.

Drewell, R. A., Lo, N., Oxley, P. R., Oldroyd, B. P. (2012). Kin conflict in insect societies: a new epigenetic perspective. Trends in Ecol & Evol, 27(7), 367–73.

Edwards, C. A., Ferguson-Smith, A. C. (2007). Mechanisms regulating imprinted genes in clusters. Curr Op in Cell Biol, 19(3), 281–289.

Elango, N., Hunt, B. G., Goodisman, M. A. D., Yi, S. V. (2009). DNA methylation is widespread and associated with differential gene expression in castes of the honeybee, Apis mellifera. PNAS, 106(27), 11206–11

Dobin, A., Davis, C. A., Schlesinger, F., Drenkow, J., Zaleski, C., Jha, S., et al. (2013). STAR: ultrafast universal RNA-seq aligner. Bioinformatics, 29(1), 15–21.

Duncan, E. J., Hyink, O., Dearden, P. K. (2016). Notch signalling mediates reproductive constraint in the adult worker honeybee. Nat Commun, 7(1), 12427.

Duncan, E. J., Leask, M. P., Dearden, P. K. (2020). Genome architecture facilitates phenotypic plasticity in the honeybee (*Apis mellifera*). Wittkopp P., editor. Molecular Biology and Evolution, 37, 1964–1978.

Fan, J., Wang, X., Xiao, R., Li, M. (2021). Detecting cell-type-specific allelic expression imbalance by integrative analysis of bulk and single-cell RNA sequencing data. Lappalainen, T., editor. PLOS Genet, 17, e1009080.

Ferguson-Smith, A. C. (2011). Genomic imprinting: The emergence of an epigenetic paradigm. Nature Reviews Genetics, 12, 565–575.

Ferreira, H. M., Alves, D. A., Cool, L., Oi, C. A., Oliveira, R. C., Wenseleers, T. (2024). Toward greater realism in inclusive fitness models: the case of caste fate conflict in insect societies. Evol Letters, qrad068.

Floc’hlay, S., Wong, E. S., Zhao, B., Viales, R. R., Thomas-Chollier, M., Thieffry, D., Garfield, D. A., & Furlong, E. E. M. (2021). Cis-acting variation is common across regulatory layers but is often buffered during embryonic development. Genome Research, 31, 211–224.

Fukushima, H. S., Takeda, H., Nakamura, R. (2023). Incomplete erasure of histone marks during epigenetic reprogramming in medaka early development. Genome Research, 33(4), 572–586.

Galbraith, D. A., Kocher, S. D., Glenn, T., Albert, I., Hunt, G. J., Strassmann, J. E., et al. (2016). Testing the kinship theory of intragenomic conflict in honey bees (*Apis mellifera*). Proc Natl Acad Sci, 113(4), 1020–5.

Galbraith, D. A., Ma, R., Grozinger, C. M. (2021). Tissue-specific transcription patterns support the kinship theory of intragenomic conflict in honey bees (*Apis mellifera*). Mol Ecol, 30(4), 1029–41.

Gardner, A., Úbeda, F. (2012). The meaning of intragenomic conflict. Nat Ecol Evol, 1(12), 1807–15.

Garrison E., Marth G. (2012). Haplotype-based variant detection from short-read sequencing. Available from: http://arxiv.org/abs/1207.3907

Gazave, E., Merqués-Bonet, T., Fernando, O., Charlesworth, B., Navarro, A. (2007). Patterns and rates of intron divergence between humans and chimpanzees. Genome Biology, 8, R21.

Gehring, M., Satyaki, P. R. (2017). Endosperm and Imprinting, Inextricably Linked. Plant Physiol, 173(1), 143–154.

Gibson, J. D., Arechavaleta-Velasco, M. E., Tsuruda, J. M., & Hunt, G. J. (2015). Biased allele expression and aggression in hybrid honeybees may be influenced by inappropriate nuclear-cytoplasmic signaling. Frontiers in Genetics, 6, 343.

Grozinger, CM. (2015). Honey bee pheromones. In J. Graham (Ed.), The hive and the honey bee. Dadant & Sons.

Haig, D. (2000). The kinship theory of genomic imprinting. Ann Rev Ecol & Sys, 31, 9–32.

Haig, D. (2008). Conflicting messages: Imprinting and internal communication. In P. d’Ettorre & D. P. Hughes (Eds.), Sociobiology of communication. Oxford University Press.

Hanna, C. W. (2020). Placental imprinting: Emerging mechanisms and functions. PLoS Genet, 16(4), e1008709.

Hanna, C. W., Kelsey, G. (2021). Features and mechanisms of canonical and noncanonical genomic imprinting. Genes & Development, 35, 821–834.

Hartfelder, K., Tiberio, G. J., Lago, D. C., Dallacqua, R. P., Bitondi, M. M. G. (2018). The ovary and its genes developmental processes underlying the establishment and function of a highly divergent reproductive system in the female castes of the honey bee, *Apis mellifera*. Apidologie, 49, 49–70.

He, X. J., Barron, A. B., Yang, L., Chen, H., He, Y. Z., Zhang, L. Z., et al. (2022). Extent and complexity of RNA processing in honey bee queen and worker caste development. iScience. 25, e104301.

Hutter, B., Helms, V., Paulsen, M. (2006). Tandem repeats in the CpG islands of imprinted genes. Genomics, 88(3), 323–32.

Hutter, B., Bieg, M, Helms, V, Paulsen, M. (2010). Imprinted genes show unique patterns of sequence conservation. BMC Genomics, 11, 649.

Inoue, A. (2023). Noncanonical imprinting: Intergenerational epigenetic inheritance mediated by Polycomb complexes. Current Opinion in Genetics & Development, 78, 102015.

Jackson, K., Robinson, G. E. (2018). Contest experience does not increase survivorship in honey bee queen duels. Insectes Sociaux, 65, 631–7.

Kanduri, C. (2016). Long noncoding RNAs: Lessons from genomic imprinting. Biochim Biophys Acta, 1859(1), 102–11.

Kent, C. F., Minaei, S., Harpur, B. A., Zayed, A. (2012). Recombination is associated with the evolution of genome structure and worker behavior in honey bees. PNAS, 109(44), 18012–7.

Kocher, S. D., Tsuruda, J. M., Gibson, J. D., Emore, C. M., ArechavaletaVelasco, M. E., Queller, D. C., Strassmann, J. E., Grozinger, C. M., Gribskov, M. R., San Miguel, P., Westerman, R., Hunt, G. J. (2015). A search for parent-of-origin effects on honey bee gene expression. G3 (Bethesda, Md.), 5, 1657–1662.

Krauss, V, Reuter, G. (2011). DNA methylation in *Drosophila* – a critical evaluation. Prog Mol Biol Transl Sci, 101, 177–91.

Kumar, D., Cinghu, S., Oldfield, A.J., Yang, P., Jothi, R. (2021). Decoding the function of bivalent chromatin in development and cancer. Genome Res, 31(12), 2170–2184.

Landt, S. G., Marinov, G. K., Kundaje, A. et al. (2012). ChIP-seq guidelines and practices of the ENCODE and modENCODE consortia. Genome Res, 22(9), 1813–1831.

Langfelder, P., Horvath, S. (2008). WGCNA: an R package for weighted correlation network analysis. BMC Bioinformatics, 9, 559.

Langmead, B., Salzberg, S. L. (2012). Fast gapped-read alignment with Bowtie 2. Nature Methods, 9, 357–359.

Lawrence, M., Huber, W., Pagès, H., Aboyoun, P., Carlson, M., Gentleman, R., et al. (2013). Software for computing and annotating genomic ranges. PLoS Comput Biol, 9(8), e1003118.

Li, H. (2013). Aligning sequence reads, clone sequences and assembly contigs with BWA-MEM. Available from: http://arxiv.org/abs/1303.3997

Love, M. I., Huber, W., Anders, S. (2014). Moderated estimation of fold change and dispersion for RNA-seq data with DESeq2. Genome Biol, 15, 550.

Lu, Y.-X., Denlinger, D. L., Xu, W.-H. Polycomb repressive complex 2 (PRC2) protein ESC regulates insect developmental timing by mediating H3K27me3 and activating prothoracicotropic hormone gene expression. J Biol Chem, 288(32), 23554–23564.

Lun, A. T. L., Smyth, G. K. (2016). Csaw: a Bioconductor package for differential binding analysis of ChIP-seq data using sliding windows. Nucleic Acids Res, 44(5), e45.

MacDonald, W.A. (2012). Epigenetic mechanisms of genomic imprinting: Common themes in the regulation of imprinted regions in mammals, plants, and insects. Genet Res Int, 585024.

Macias-Valesco, J. F., Pierre, C. L. St., Wayhart, J. P., Yin, L., Spears, L., et al. (2022). Parent-of-origin effects propagate through networks to shape metabolic traits. eLife, 11, e72989.

Makert, G. R., Paxton, R. J., Hartfelder, K. (2006). Ovariole number—a predictor of differential reproductive success among worker subfamilies in queenless honeybee (*Apis mellifera L.*) colonies. Behavioral Ecology and Sociobiology, 60, 815–825.

Maleszka, R., Kucharski, R. (2022). Without mechanisms, theories and models in insect epigenetics remain a black box. Trends in Genetics, 38(11), P1108–11.

Martinez, M. E., Cox, D. F., Youth, B. P., Hernandez, A. (2016). Genomic imprinting of *DIO3*, a candidate gene for the syndrome associated with human uniparental disomy of chromosome 14. European J of Hum Gen, 24, 1617–1621.

Matsuura, K. (2019). Genomic imprinting and evolution of insect societies. Population Ecology, 62(1), 38–52.

McEwen, K. R., Ferguson-Smith, A. C. (2010). Distinguishing epigenetic marks of developmental and imprinting regulation. Epigenetics & Chromatin, 3, 2.

McKenna, A., Hanna, M., Banks, E., Sivachenko, A., Cibulskis, K., Kernytsky, A., Garimella, K., Altshuler, D., Gabriel, S., Daly, M., et al. (2010). The Genome Analysis Toolkit: A MapReduce framework for analyzing next-generation DNA sequencing data. Genome Res, 20, 1297–1303.

Obregón, F. P., Soto, P., Lavín, J. L., Cortázar, A. R., Barrio, R., Aransay, A. M., Cantera, R. (2018). Cluster Locator, online analysis and visualization of gene clustering. Bioinformatics, 34(19), 3377–3379.

Oldroyd, B. P., Yagound, B. (2021). Parent-of-origin effects, allele-specific expression, genomic imprinting, and paternal manipulation in social insects. Phil Trans Royal Soc London B Biol Sci, 376(1826), 20200425.

Olney, K.C., Gibson, J.D., Natri, H.M., Underwood, A., Gadau, J., Wilson, M.A. (2021). Lack of parent-of-origin effects in *Nasonia* jewel wasp: A replication and extension study. PLoS One, 16(6), e0252457.

Page, R. E. (2013). The spirit of the hive. Harvard University Press.

Patten, M. M., Ross, L., Curley, J. P., Queller, D. C., Bonduriansky, R., Wolf, J. B. (2014). The evolution of genomic imprinting: Theories, predictions and empirical tests. Heredity, 113, 119–128.

Patten, M. M., Cowley, M., Oakey, R. J., Feil, R. (2016). Regulatory links between imprinted genes: Evolutionary predictions and consequences. Proceedings of the Royal Society B: Biological Sciences, 283, 20152760.

Pegoraro, M., Marshall, H., Lonsdale, Z. N., Mallon, E. B. (2017). Do social insects support Haig’s kin theory for the evolution of genomic imprinting? Epigenetics, 12, 725–742.

Pertea, G., Pertea, M. (2020). GFF Utilities: GffRead and GffCompare. F1000Res, 9, ISCB Comm J-304.

Pigliucci, M. (2001). Phenotypic plasticity: Beyond nature and nurture. Johns Hopkins University Press.

Prachumwat, A., DeVincentis, L., Palopoli, M. F. (2004). Intron size correlates positively with recombination rate in *Caenorhabditis elegans*. Genetics, 166(3), 1585–90.

Queller, D. C. (2003). Theory of genomic imprinting conflict in social insects. BMC Evolutionary Biology, 3, 15.

Quinlan, A. R., Hall, I. M. (2010). BEDTools: a flexible suite of utilities for comparing genomic features. Bioinformatics, 26, 841–84.

R Core Team. (2024). R: A language and environment for statistical computing. R Foundation for Statistical Computing, Vienna, Austria. https://www.R-project.org/.

Radford, E. J., Ferrón, S. R., Ferguson-Smith, A. C. (2011). Genomic imprinting as an adaptive model of developmental plasticity. FEBS Letters, 2059–2066.

Raas, M. W. D., Zijlmans, D. W., Vermeulen, M., Marks, H. (2022). There is another: H3K27me3-mediated genomic imprinting. Trends Genet, 38(1), 82–96.

Robinson, M. D., McCarthy, D. J., Smyth, G. K. (2010). edgeR: a Bioconductor package for differential expression analysis of digital gene expression data. Bioinformatics, 26(1), 139–140.

Roy, M., Kim, N., Xing, Y., Lee, C. (2008). The effect of intron length on exon creation ratios during the evolution of mammalian genomes. RNA, 14(11), 2261–2273.

Sagili, R. R., Metz, B. N., Lucas, H. M., Chakrabarti, P., Breece, C. R. (2018). Honey bees consider larval nutritional status rather than genetic relatedness when selecting larvae for emergency queen rearing. Scientific Reports, 8, 7679.

Sandovici, I., Kassovska-Bratinova, S., Vaughan, J. E., Stewart, R., Leppert, M., Sapienza, C. (2006). Human imprinted chromosomal regions are historical hot-spots of recombination. PLoS Genet, 2(7), e101.

Sanli, I., Feil, R. (2015). Chromatin mechanisms in the developmental control of imprinted gene expression. Int J Biochem & Cell Biol, 67, 139–147.

Sherman, B. T., Hao, M., Qiu, J., Jiao, X., Baseler, M.W., Lane, H.C., Imamichi, T., Chang, W. (2022). DAVID: a web server for functional enrichment analysis and functional annotation of gene lists (2021 update). Nucleic Acids Res, 50, W216–W221.

Slater, P. G., Dapper, A. L., Harpur, B. A. (2022). Haploid and sexual selection shape the rate of evolution of genes across the honey bee (*Apis mellifera* L.) genome. Genome Biol & Evol, 14(6), evac063.

Smit, A. F. A., Hubley, R., Green, P. (2015). RepeatMasker Open-4.0. Available from: http://www.repeatmasker.org

Smith, N. M. A., Yagound, B., Remnant, E. J., Foster, C. S. P., Buchmann, G., Allsopp, M. H., Kent, C. F., Zayed, A., Rose, S. A., Lo, K., Ashe, A., Harpur, B. A., Beekman, M., Oldroyd, B. P. (2020). Paternally-biased agene expression follows kin-selected predictions in female honey bee embryos. Molecular Ecology, 29, 1523–1533.

Soliman, H. K., Coughlan, J. M. (2024). United by conflict: Convergent signatures of parental conflict in angiosperms and placental mammals. *J Heredity*, esae009.

Soneson, C., Love, M. I., Robinson, M. (2022). Differential analyses for RNA-seq: transcript-level estimates improve gene-level inferences. F1000Research, 4.

Storer, B. E., Kim, C. (1990). Exact Properties of Some Exact Test Statistics for Comparing Two Binomial Proportions. J. Am. Stat. Assoc, 85, 146–155.

Tan, J., Yang, X., Zhuang, L., Jiang, X., Chen, W., Lee, P. L., Karuturi, R. M., Tan, P. B. O., Liu, E. T., Yu, Q. (2007). Pharmacologic disruption of Polycomb-repressive complex 2-mediated gene repression selectively induces apoptosis in cancer cells. Genes Dev, 21, 1050–1063.

Thompson, G. J., Chernyshova, A. M. (2021). Caste Differentiation: Genetic and Epigenetic Factors. In C. K. Starr (Ed.), Encyclopedia of social insects (pp. 165–176). essay, Springer International Publishing.

van Ekelenburg, Y. S., Hornslien, K. S., Hautegem, T. V., Fendrych, M., Isterdael, G. V., et al. (2022). Spatial and temporal regulation of parent-of-origin allelic expression in endosperm. Plant Physiol, 191(2), 986–1001.

Wallberg, A., Bunikis, I., Pettersson, O. V., Mosbech, M.-B., Childers, A. K., Evans, J. D., Mikheyev, A. S., Robertson, H. M., Robinson, G. E., Webster, M. T. (2019). A hybrid de novo genome assembly of the honeybee, *Apis mellifera*, with chromosome-length scaffolds. BMC Genomics, 20, 275.

Wang, X., Clark, A. G. (2014). Using next-generation RNA sequencing to identify imprinted genes. Heredity, 113, 156–166.

Wang, M., Xiao, Y., Li, Y., Wang, X., Qi, S., Wang, Y., Zhao, L., et al. (2021). RNA m6A modification functions in larval development and caste differentiation in honeybee (*Apis mellifera*). Cell Reports, 34(1), e108580.

Wang, Q., Li, M., Wu, T., Zhan, L., Li, L., Chen, M., et al. (2022). Exploring epigenomic datasets by ChIPseeker. Current Protocols, 2(10), e585.

Wojciechowski, M., Lowe, R., Maleszka, J., Conn, D., Maleszka, R., Hurd, P. J. (2018). Phenotypically distinct female castes in honey bees are defined by alternative chromatin states during larval development. Genome Res, 28(10), 1532–1542.

Wolf, J. B. (2013). Evolution of genomic imprinting as a coordinator of coadapted gene expression. PNAS, 110(13), 5085–90.

Wight, M., Werner, A. (2013). The functions of natural antisense transcripts. Essays Biochem, 54, 91–101.

Wu X., Galbraith D.A., Chatterjee P., Jeong H., Grozinger C.M., Yi S.V. (2020). Lineage and parent-of-origin effects in DNA methylation of Honey Bees (*Apis mellifera*) revealed by reciprocal crosses and whole-genome bisulfite sequencing. Genome Biol Evol, 12(8), 1482–92.

Xia, W., Xie, W. (2020). Rebooting the epigenomes during mammalian early embryogenesis. Stem Cell Reports, 15(6), 1158–1175.

Yang, Y., Zhao, T., Li, Z., Qian, W., Peng, J., Wei, L, et al. (2021). Histone H3K27 methylation-mediated repression of *Hairy* regulates insect developmental transition by modulating ecdysone biosynthesis. PNAS, 118(35), e2101442118.

Yoon, K. J., Cunningham, C. B., Bretman, A., Duncan, E. J. (2023). One genome, multiple phenotypes: decoding the evolution and mechanisms of environmentally induced developmental plasticity in insects. Biochem Soc Trans, 51(2), 675–689.

Young, M. D., Willson, T. A., Wakefield, M. J., Trounson, E., Hilton, D. J., Blewitt, M. E., Oshlack, A., Majewski, I. J. (2011). ChIP-seq analysis reveals distinct H3K27me3 profiles that correlate with transcriptional activity. Nucleic Acids Res, 39(17), 7415–7427.

Zhang, Y., Meyer, C. A., Eeckhoute, J., Johnson, D. S., Bernstein, B. E., et al. (2008). Model-based analysis of ChIP-seq (MACS). Genome Biology, 9, R137.

Zhang, Y., Li, Z., He, X., Wang, Z., Zeng, Z. (2023). H3K4me1 modification functions in caste differentiation in honey bees. Int J Mol Sci, 24(7), 6217.

Zeng, J., Yi, S. V. (2010). DNA methylation and genome evolution in honeybee: Gene length, expression, functional enrichment covary with the evolutionary signature of DNA methylation. Genome Biol Evol, 2: 770–80.

Zhu, A., Ibrahim, J. G., Love, M. I. (2019). Heavy-tailed prior distributions for sequence count data: removing the noise and preserving large differences. Bioinformatics, 35(12), 2084–2092.

Zhu, L., Zhang, Y., Zhang, W., Yang, S., Chen, Y.-Q., Tian, D. (2009). Patterns of exon-intron architecture variation of genes in eukaryotic genomes. BMC Genomics, 10, 47.

